# MHC restriction is structurally encoded by the T cell receptor germline

**DOI:** 10.64898/2026.09.24.754241

**Authors:** Nan Wang, Marta T. Borowska, Kevin M. Jude, Xiaojing Chen, Chunyu Wang, Xinyu Xiang, Liu Daisy Liu, Rong Ma, Anshika Motiani, Steven C. Wilson, Wanlu Liu, K. Christopher Garcia

## Abstract

MHC restriction — the requirement for αβ T cell receptors (TCRs) to recognize Major Histocompatibility Complex (MHC) molecules when presenting peptide antigens — is a pillar of adaptive immunity. Here, we provide direct structural evidence for TCR germline-encoded recognition of MHC molecules irrespective of antigenic peptide. We engineered “germline-like” TCRs to eliminate CDR3 specificity for peptide, and a molecular clamping strategy to trap ultra-low-affinity complexes with peptide-MHC for cryo-EM. We find that germline-like TCR/pMHC docking modes are identical to wild-type T cell receptors with intact CDR3 loops, across distinct peptide antigens and MHC alleles. Analysis of human TCR repertoires reveals enrichment of specific V genes and Vα-Vβ pairing patterns associated with HLA-A*02 contexts. Thus, the TCR germline encodes the blueprint for MHC restriction, providing a sequence-based framework that could inform computational and AI-driven prediction of TCR specificity.

## Introduction

Adaptive immunity relies on lymphocytes that can recognize an essentially unlimited universe of molecular threats, each through a distinct surface receptor. Among our three lineages of antigen receptors, αβ T cell receptors (TCRs) are unique in their requirement to recognize protein antigens as processed peptides presented by Major Histocompatibility Complex (MHC) molecules, a phenomenon termed MHC restriction ^1,2^. Unlike antibodies, which are not limited to seeing any particular structural class of proteins or ligands, the TCR is restricted to engagement of a composite surface composed of both the α-helices bracketing the MHC binding groove and the exposed residues of the bound peptide ^3–6^. The TCR, while incredibly diverse, is generally considered to have a ‘bias’ for binding to MHC molecules ^7–9^. This bias is what allows the T cells to successfully undergo positive selection and function in an MHC-restricted manner ^10–12^.

The forces that shape MHC restriction by the TCR have been studied extensively without clear resolution ^13^, but have been explained by two contrasting theories. The co-evolution model is historically rooted in early Jerne hypotheses of antigen receptor V-gene germline-encoded reactivity with histocompatibility antigens ^14^. This theory, which predated the discovery of the T cell receptor, opined that antigen receptors on cells and MHC molecules co-evolved over time, leading to features within the TCR that are intrinsically compatible with MHC ^15–17^. Supporting this, genetic and structural analyses of TCR-peptide-MHC complexes show that the more conserved, germline encoded parts of the TCR (the CDR1 and CDR2 loops) tend to interact directly with the conserved α-helical regions of the MHC molecule (**Figures 1A and 1B**) ^4,18–20^. This suggests a built-in “fit” that predisposes the TCR to engage with the MHC protein ^21,22^. On the other hand, germline components of the TCR can also form peptide contacts, and CDR3s contact the MHC helices, so the division of labor is not absolute ^23^.

**Figure 1.**
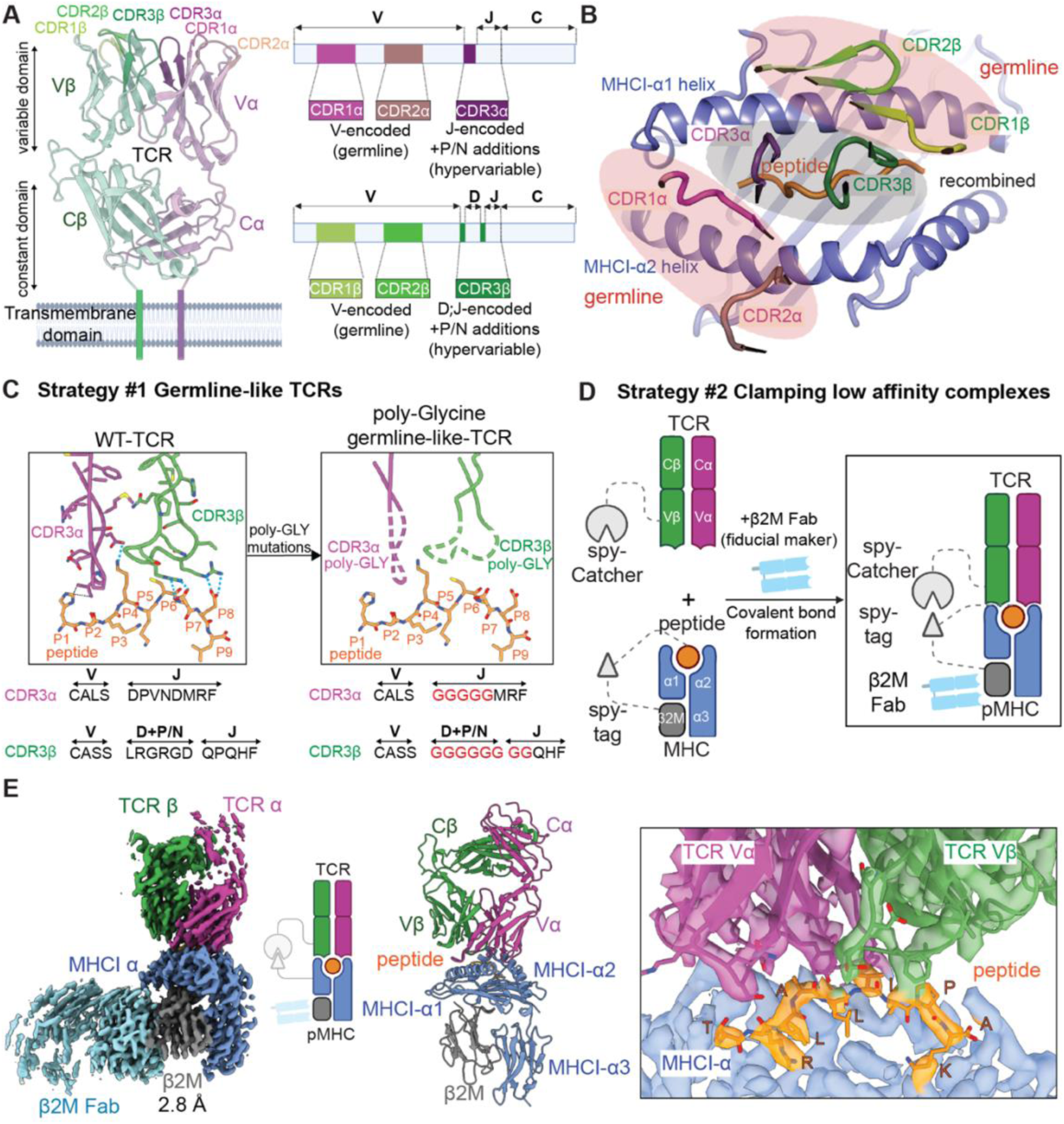
Strategies for trapping ultra-low affinity germline-like TCR-pMHC for structure determination. (A) Overview of TCR architecture and gene organization. Left: structural arrangement of TCR α and β chains. Right: schematic of the corresponding gene segments showing that CDR1 and CDR2 loops are germline-encoded, whereas CDR3 loops arise from V(D)J recombination with N/P nucleotide additions. (B) Canonical topology of the TCR-pMHC interface showing the diagonal orientation of the TCR variable domains across the MHC α1 and α2 helices. Germline-encoded CDR1 and CDR2 loops contact the MHC surface, whereas CDR3 loops engage the bound peptide. (C) Design of germline-like TCRs. In the wild-type (WT) TCR, both CDR3α and CDR3β loops interact directly with the peptide (left, representative structure PDB 4QRP). In germline-like-engineered constructs, these loops are replaced by poly-glycine linkers (right), removing peptide-specific contacts while preserving germline CDR1/CDR2 frameworks for MHC recognition. (D) Clamping strategy for stabilizing low-affinity germline-like TCR-pMHC complexes. A SpyTag on the pMHC forms a covalent isopeptide bond with a SpyCatcher fused to the TCR β chain, while a β2M-specific Fab provides a fiducial marker for cryo-EM alignment. (E) Representative cryo-EM reconstruction of the AS4.2-PRPF3-HLA-B*27 complex using clamping strategy at 2.8 Å resolution. Left: overall map and fitted model showing TCR α (purple), TCR β (green), MHC (blue), β2M (gray) and β2M Fab (cyan). Right: zoomed-in view highlighting the peptide (orange) within the groove and its interaction with the TCR α and β chains.

In contrast to the coevolution model, other models posit that CDR3 recombination in response to peptide antigens drives MHC restriction, or that the bias is not intrinsic but is largely extrinsically imposed by the selection process in the thymus and the necessity of co-receptors (CD4 or CD8). CD4 (for MHC Class II) and CD8 (for MHC Class I) co-receptors, by virtue of their association with the lymphocyte-specific protein tyrosine kinase (Lck), are essential for T cell signaling ^21,24^. This co-receptor requirement focuses the repertoire toward MHC-reacting TCRs, giving the appearance of intrinsic TCR-MHC affinity when in fact there may not be.

Despite decades of study ^19,22,25^, unambiguous structural evidence for intrinsic, peptide-independent TCR-MHC affinity has been lacking. The fundamental obstacle is that, in all wild-type TCR-pMHC structures solved to date, the receptor contains intact CDR3 loops and is determined in the presence of a specific peptide. As CDR3 contributes to both MHC and peptide contacts, and CDR1/CDR2 loops can also touch peptide side chains, it has not been possible to isolate the contribution of the germline V-domains from that of the adaptive CDR3 in any single structure. What is needed is a structural approach that retains the germline framework intact while genuinely abolishing CDR3-mediated contacts. Resolving this classical debate carries practical weight as well: if MHC engagement is genuinely hard-coded in the germline rather than an emergent property of selection, it constitutes a fixed structural constraint that could substantially narrow the sequence space that computational and AI-based approaches must search when predicting TCR specificity for peptide-MHC.

Here, we provide direct structural evidence that TCR germline V-regions bind MHC independently of CDR3 and peptide interactions. Using “germline-like” TCRs devoid of CDR3 peptide-specificity and a molecular clamping strategy to trap ultra-low-affinity complexes, we find that germline-like TCR/pMHC docking geometries are virtually identical to wild-type peptide-specific TCR/pMHC complexes across distinct peptides and MHC alleles. These findings demonstrate that MHC restriction is encoded in the TCR germline, and that the rules governing MHC engagement should in principle be predictable from TCR V-region sequence alone, a prerequisite for sequence-based computational prediction of TCR specificity.

## Results

### Clamping strategies to capture low-affinity TCR-pMHC complexes

To capture germline-mediated TCR-MHC complexes, we devised a two-part engineering strategy. First, to eliminate peptide-specific CDR3 contacts while preserving the germline CDR1/CDR2 framework, we replaced both CDR3α and CDR3β with poly-glycine linkers (**Figure 1C**). Glycine residues carry no side chains and are conformationally flexible, so the poly-glycine stretches are incapable of sequence-specific contacts with either peptide or MHC, yet do not destabilize the overall receptor fold. (**Figure 1C**). Second, because germline-like TCR-pMHC complexes are predicted to be too ultra-low affinity and transient to persist in cryo-EM samples, we developed a “clamping” approach (**Figure 1D**).

We first wished to test the efficiency of complex formation of low affinity versus high affinity TCR-pMHC complexes by cryoEM in absence of clamping. We compared a human TCR that recognized the PAP tumor antigen ^26^ to “higher-affinity” complex of TCR A3A specific for MAGE-A3-HLA-A*01. We first assessed a direct binding affinity between TCR156 specific for PAP-HLA-A*02 (*K*_D_ of 47.4 μM) and compared it with “higher-affinity” complex of TCR A3A specific for MAGE-A3-HLA-A*01 (*K*_D_ of 1.8 μM) (**Figures S1A and S1B**). 2D classification of the lower-affinity TCR156-PAP-HLA-A*02 complex shows a low proportion of assembled particles under these conditions, indicating limited complex formation. By comparison, the higher-affinity A3A-MAGE-A3-HLA-A*01 complex (*K*_D_ ∼1.8 μM) showed clear complex formation by 2D classification (**Figures S1D and S1E**). These results underscore the technical challenge of capturing weak or transient complexes for structural studies, especially those involving germline-like TCRs.

To overcome this limitation, we developed a clamping strategy to enable cryo-EM analysis of complexes that would otherwise be too transient for visualization. We used a SpyTag-SpyCatcher linkage to physically tether TCR and pMHC, coupled with a Fab fragment for fiducial imaging alignment (**Figure 1D**) ^27^. The goal of the clamping strategy was to raise the effective concentrations of the low affinity TCR and pMHC sufficiently to form complex, while permitting enough orientational freedom that the molecules were not constrained to binding in artifactual ways. In our clamping constructs, SpyCatcher was fused to the N terminus of TCRβ through a flexible 13-amino acid Gly/Ser linker, and the 13-amino acid SpyTag was inserted between the peptide and β₂m within the single-chain pMHC, flanked by a (G₃S)₃ linker on the peptide side and an additional 8-residue GS linker before β₂M. These long linkers provided ample orientational freedom to allow for a wide range of docking geometries (**Figure S2**). Using this clamping method, we determined a cryo-EM structure with an existing X-ray TCR-pMHC structure AS4.2-PRPF3-HLA-B*27 (**Figures 1E and S3**) ^28^. The cryo-EM reconstructions displayed close agreement with available crystal structures, after alignment on the HLA-B27 α1-α2 platform, the cryo-EM and crystal structures showed a similar global docking geometry (TCR V-domain Cα RMSD = 2.22 Å over 229 atom pairs; crossing angle 49.47° vs 44.69°, Δ = 4.77°), indicating that the clamping strategy can faithfully capture low-affinity or transient interactions without introducing detectable structural artifacts (**Figure S4**). The resulting maps reached resolutions to 2.8 Å, sufficient to resolve interfacial features such as CDR-MHC contacts.

With the clamping strategy validated, we then asked if a clamped germline-like TCR-pMHC complex could be visualized by cryo-EM. We generated a ‘germline-like’ TCR156, by replacing the CDR3 loops by mutating TCR CDR3α residues 92-95 (NNAR) to GGGG and TCR CDR3β residues 95-102 (VAGSPEAF) to GGGGGGGG (TCR156 numbering). After replacing the CDR3α and CDR3β loops, the germline-like version of TCR156 bound PAP-HLA-A*02 only very weakly, with an estimated *K*_D_ of hundreds of uM, at the lower sensitivity limit of SPR (**Figure S1C**).

### Germline-like TCRs adopt similar docking geometry as wild-type TCRs

We next applied the clamping strategy to determine the cryo-EM structures of germline-like versions of TCR156 (TRAV12-2, TRBV9) in complex with PAP-HLA-A*02 (**Figure 2A**). The germline-like TCR156 maintained a conserved global docking geometry relative to its wild-type counterpart structure (**Figures 2B, 2C and S5A**) ^29^. The poly-glycine CDR3 loops were largely invisible in the map, consistent with their flexibility and lack of engagement with the pMHC (**Figure S6A**), suggesting their minimal contribution to its docking configuration.

**Figure 2.**
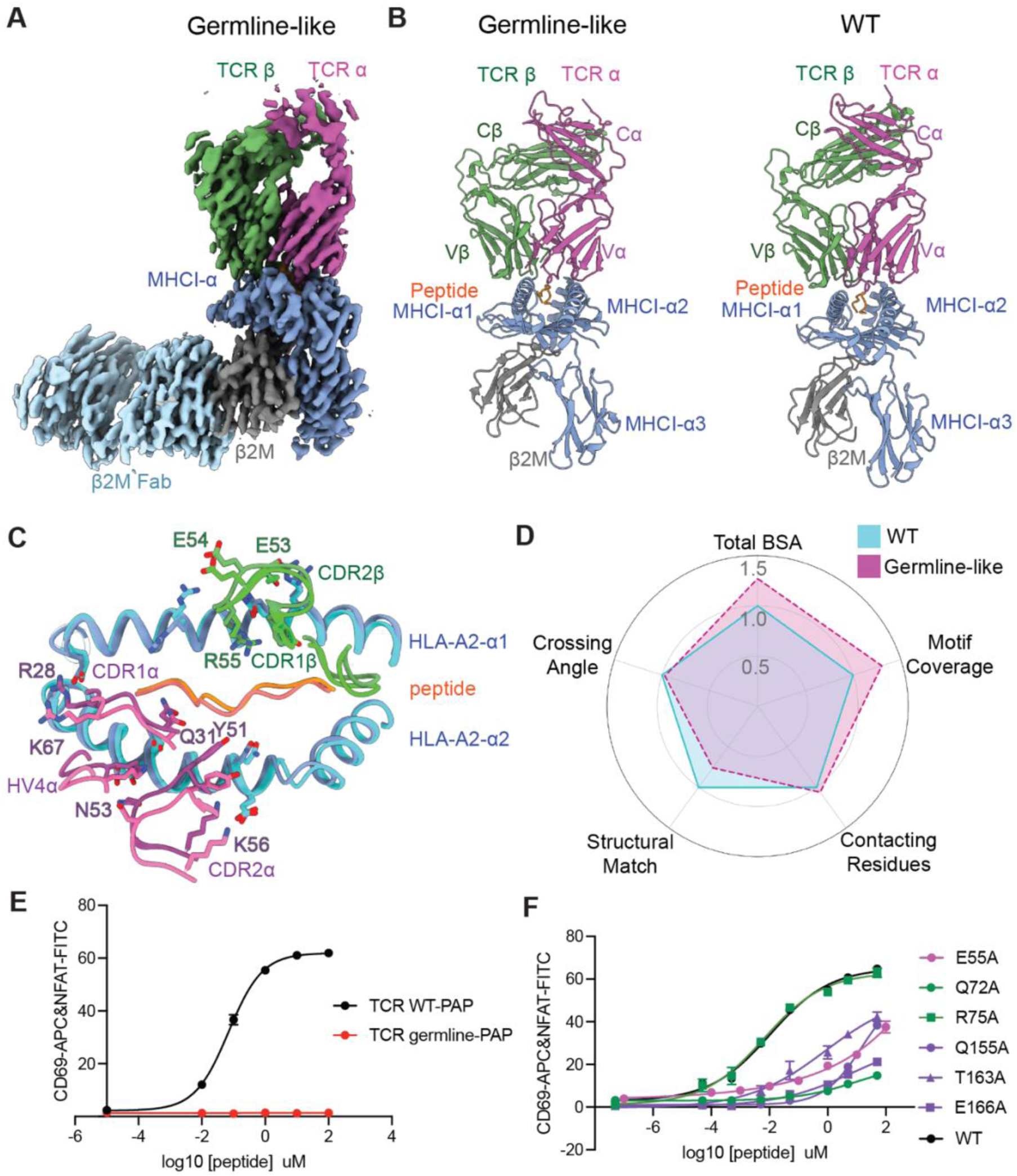
Germline-like TCR156 adopts similar docking geometry with HLA-A*02 as wild-type TCR156. (A) Cryo-EM reconstruction of the germline-like TCR156-PAP-HLA-A*02 complex stabilized using the clamping strategy, showing TCR α (magenta), TCR β (green), MHC (blue), and β2M Fab (gray). (B) Structural comparison between germline-like (left) and wild-type (WT, right) TCR156 bound to PAP-HLA-A*02. Despite the replacement of both CDR3α and CDR3β loops with poly-glycine linkers, the germline-like complex maintains similar overall docking geometry and diagonal orientation as the WT TCR. (C) Top-down view of the superimposed interface between the WT (light colors) and germline-like (darker colors) TCR-pMHC complexes. Key interacting residues in the germline-encoded CDR1 and CDR2 loops are represented as sticks. MHC residues are numbered according to the standard HLA-A*02 mature polypeptide sequence, independent of their positions within the single-chain trimer (SCT) construct. (D) Multi-dimensional comparison of WT and germline-like TCR-pMHC interaction. A radar plot quantifying five biophysical and structural metrics of the germline-like TCR-pMHC interface (magenta dashed line) normalized to the WT values (cyan solid line, set at 1.0). The metrics include the variable domain RMSD (1.176 Å), the number of contacting residues (18 for germline-like vs. 17 for WT), the total buried surface area (BSA = 1025.5 Å^2^ for germline-like vs. 806.6 Å^2^ for WT), the motif coverage index (1.30 for germline-like), and the total solvent accessible surface area (SASA; 34130 for germline-like vs. 33655 for WT). (E-F) Peptide-induced activation of Jurkat reporter cells expressing WT or germline TCR156 (E). Activation assays using alanine-substituted HLA-A*02 variants expressed on K562 cells show the corresponding changes in CD69/NFAT readouts (F).

To quantitatively evaluate this conserved interface, we utilized a multi-dimensional radar plot to compare several key structural and biophysical metrics (**Figure 2D**). The two complexes exhibited remarkable similarity across all parameters. First, the variable domains showed high structural congruence with a minimal displacement (Cα RMSD) of 1.18 Å. Second, the “crossing angle”, which defines how the TCR sits across the MHC, was nearly identical (64.26° for WT versus 62.34° for germline). Third, the total surface area buried upon binding (BSA) remained high (1025.5 Å^2^ for germline vs. 806.6 Å^2^ for WT), supported by a consistent number of physical contacts (18 for germline vs. 17 for WT). Finally, the “motif coverage index,” which measures the preservation of core interacting residues, was 1.30 relative to WT. Together, these data demonstrate that the germline-encoded framework provides a stable and consistent structural “blueprint” that anchors the TCR in a canonical docking mode independently of the CDR3 loops.

The germline-mediated interface is characterized by a stable network of conserved interactions between the TCR V-region loops (CDR1/2) and the MHC α-helices (**Figures 2C and S5B**). In the α-chain interface, a cluster of polar residues in CDR1α (Arg28) and CDR2α (Tyr51, Asn53) forms a recurrent contact motif with the MHC α2-helix (Gln155, Glu166). This framework is reinforced by HV4α, where Lys67 acts as a bridge to the upper portion of the α2-helix. Similarly, the CDR2β loop anchors onto the MHC α1-helix through a complementary set of hydrogen bonds (e.g., Glu53^β^-Arg75^α^^1^ and Arg55^β^-Gln72^α^^1^). Interestingly, Gln31α, positioned at the base of the CDR1α loop, extended deep into the peptide-binding groove to contact the main-chain atoms of the peptide (Leu2-Ser4) via backbone-level hydrogen bonds. This interaction likely senses peptide conformation without contributing to side-chain specificity, providing a structural rationale for how the germline framework recognizes the MHC-peptide complex in a largely peptide-independent yet geometrically consistent manner.

We examined whether the germline-mediated contacts observed in the structure are necessary for functional T cell activation. Jurkat NFAT reporter cells expressing wild-type (WT) TCR156 were robustly activated by antigen-presenting cells pulsed with the cognate PAP peptide (TLMSAMTNL), showing strong upregulation of CD69-APC and NFAT-FITC signals in a peptide dose-dependent manner (**Figure 2E**). In contrast, germline-like TCR156 failed to elicit any detectable activation across all peptide concentrations, consistent with its extremely weak binding affinity in SPR assays. These results indicate that CDR3-peptide interactions are essential for productive signaling.

To dissect if the germline-encoded contacts are required for signaling in the WT TCR interaction with HLA-A*02-PAP, we introduced alanine substitutions at key interfacial residues on HLA-A*02 helices identified from the cryo-EM structures (**Figure 2F**). Mutations in Arg75, located in peripheral positions of the interface, had minimal effect on activation, whereas substitutions at Gln72 on MHC α1 helix, which form part of the conserved germline contact network with the TCR156 CDR2β, caused dramatic reductions in signaling potency. Similarly, substitutions of Gln155, Thr163, and Glu166 on the α2 helix, residues that engage CDR2α and HV4α, caused progressive losses in signaling potency. Substitution of Glu55 on the α1 helix, which contacts CDR1α produced a modest reduction in signaling potency. These functional data mirror our structural observations, suggesting that MHC restriction relies on a distributed but cooperative network of germline-mediated interactions, where the collective integrity of these ’motifs’ is essential for efficient receptor triggering.

### Germline-like TCRs engage MHC in a shared, peptide-independent geometry

To interrogate how the germline-mediated docking mode changes with different peptide sequences, we solved additional cryo-EM structures of germline-like TCR156 bound to HLA-A*02 presenting three peptides completely distinct from PAP (**Figures S7 and S8**). The resulting reconstructions for PAP (GL156-PAP-A2), MART1 (GL156-MART1-A2), NYESO-1 (GL156-NYESO-A2), and the poly-alanine variant (GL156-polyA-A2) reached overall resolutions of 2.9, 3.1, 3.0, and 3.2 Å, respectively. The peptide panel included MART1 (ELAGIGILTV), NYESO-1 (SLLMWITQV), and a poly-alanine variant (ALAAAAAAL), in which only the canonical P2 and P9 anchor residues were retained to ensure stable MHC binding while minimizing potential interactions between the TCR and peptide side chains (**Figure 3A**). These peptides differ substantially in both sequence and length, i.e., PAP, NYESO-1, and poly-Ala being 9-mers, whereas MART1 is a 10-mer, therefore providing a stringent test of peptide dependence.

**Figure 3.**
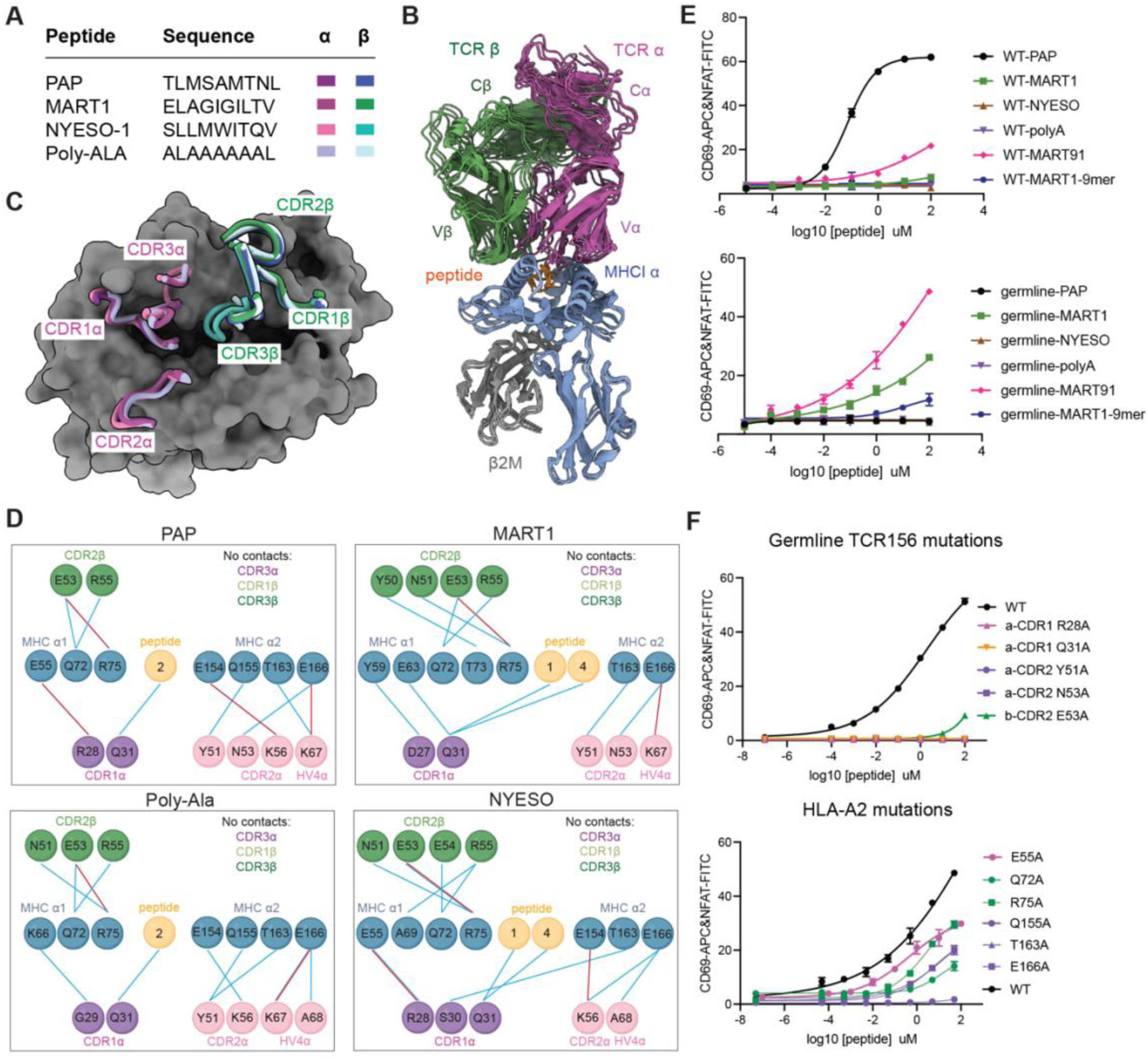
Germline-like TCR156 engage HLA-A2 in a shared, peptide-independent geometry. (A) Peptides used in structural and functional analyses. PAP, MART1, NYESO-1 and a poly-alanine variant (poly-Ala) are shown with their sequences; colored bars indicate the peptides used in the α- and β-chain structures in panels. (B) Superimposed cryo-EM structures of germline-like TCR156 bound to HLA-A*02 presenting different peptides (PAP, MART1, NYESO-1, and poly-Ala), showing TCR α (magenta), TCR β (green), MHC (yellow), β2M (gray), and peptides (orange). (C) View down the peptide-MHC groove highlighting the footprints of the CDR loops on HLA-A*02. CDR1/2α and CDR1/2β (magenta and green) sit over the α2 and α1 helices, respectively, whereas the CDR3 loops are positioned away from the peptide surface. (D) Schematic representation of germline contacts for germline-like TCR156 bound to PAP, MART1, NYESO-1 and poly-Ala. Blue lines indicate H-bonds within 3.6 Å distance and red lines indicate salt bridges, both as predicted with PDBePISA. For each complex, CDR1/2α (magenta) predominantly contact the MHC α2 helix, CDR1/2β (green) contact the α1 helix, and the peptide (orange) contributes no side chain interactions. MHC residues are numbered according to the standard HLA-A*02 mature polypeptide sequence, independent of their positions within the single-chain trimer (SCT) construct. (E) Peptide-dose response of WT (top) and germline-like (bottom) TCR156 Jurkat reporter cells stimulated with T2 cells pulsed with PAP, MART1, NYESO-1, poly-Ala, the cross-reactive bacterial peptide MART1-91 and a MART1 9-mer variant, measured by CD69/NFAT reporter readout. PAP serves as the reference ligand. (F) Functional impact of germline-mediated contacts. Top, CD69/NFAT responses of Jurkat cells expressing germline-like TCR156 variants with alanine substitutions in CDRα or CDRβ, stimulated with MART1-91 pulsed T2 cells. Bottom, responses of germline-like TCR156 to HLA-A*02 mutants carrying alanine substitutions in HLA-A*02 helix residues, expressed on K562 cells and pulsed with MART1-91.

The cryo-EM reconstructions show that germline-like TCR156 adopted a shared docking geometry across all peptide contexts (**Figure 3B**). Superimposition of the complexes revealed that the core docking orientation is highly preserved, with an average RMSD of ∼2.6 Å (ranging from 2.0 to 3.7 Å) for the TCR variable domains when aligned on the MHC α-helices. This degree of structural conservation is remarkable given the low-affinity nature of the germline-MHC interaction. The overall docking polarity remained stable, with crossing angles clustered within a narrow range of ∼61° to 72° (**Figure 3B**). Notably, even in the poly-Ala complex, where nearly all potential peptide-TCR side chain contacts were eliminated, the positioning of the CDR1 and CDR2 loops over the MHC α-helices remained unperturbed. Comparison of the aligned structures revealed that the CDR3 loops occupied similar positions relative to the MHC surface in all complexes (**Figure 3C**). In several germline structures, the CDR3α loop appeared highly flexible, consistent with its lack of engagement to pMHC (**Figure S6**). No direct peptide-CDR3 interactions were detected in some germline complexes.

Detailed inspection of the interface revealed that CDR1 and CDR2 residues form a recurring network of hydrogen bond contacts with the MHC α-helices, preserved in all four structures (**Figures 3D and S9**). These shared interfacial contacts, particularly between CDR1/2 and the MHC α helix, anchor the receptor in a common orientation regardless of the bound peptide. Notably, one germline residue, Gln31α, consistently formed backbone-level hydrogen bonds with the peptide main chain (**Figure 3D**). This interaction was preserved across both 9-mer (PAP, NYESO-1, poly-Ala) and 10-mer (MART1) peptides, adjusting subtly to accommodate peptide length while maintaining its hydrogen-bond geometry. The persistence of this Gln31-mediated contact highlights that while germline-like TCRs can engage MHC independently of peptide side chains, they nevertheless sense the peptide backbone conformation through shared framework residues.

We compared PAP, MART-1, NYESO-1, and a poly-alanine variant for their ability to activate WT or germline-like TCR156 T cells. In co-culture assays, PAP efficiently activated WT TCR156 but failed to stimulate the germline-like receptor (**Figure 3E**). Neither WT nor germline-like TCR156 responded to NYESO-1 or the poly-alanine peptide. By contrast, germline-like TCR156 displayed a weak but reproducible activation in response to MART1, reaching ∼27% and ∼10% CD69⁺ NFAT⁺ cells with the MART1 10-mer and 9-mer (AAGIGILTV), respectively, whereas WT TCR156 remained largely unresponsive (**Figure 3E**). To further enhance the activation signal, we tested MART1-91, a Mycobacterium-derived peptide that is cross-reactive with MART1 ^30^. MART1-91 elicited a substantially stronger response (∼62% CD69⁺ NFAT⁺) from germline-like TCR156 despite the absence of sequence-specific CDR3-mediated peptide contacts, while WT TCR156 remained a modest activation (∼22%) (**Figures 3E and S10**). These findings suggest that the poly-Glycine CDR3 loops formed adventitious interactions with the MART1-91 peptide that supplemented the germline-MHC contacts to support weak but productive signaling when the overall docking geometry is favorable.

The enhanced signaling activity of germline-like TCR156 to MART1-91 presented an opportunity to re-examine the importance of the germline-MHC contacts in signaling in the context of the germline-like TCR 156. We performed alanine scanning mutagenesis of key germline-facing residues on the TCR α-chain, including CDR1α-R28A, CDR1α-Q31A, CDR2α-Y51A, and CDR2α-N53A, each of which completely abolished T cell activation. Mutation of the β-chain contact CDR2β-E53A attenuated activation rather than eliminating it, indicating partial contribution (**Figure 3F, top**). In contrast, reciprocal alanine substitutions on HLA-A*02 (E55A, Q72A, R75A, Q155A, T163A, and E166A) resulted in graded reductions in signaling rather than complete loss (**Figure 3F, bottom**), confirming our earlier mutational scan of the wild-type TCR156, and showing that the TCR-pMHC interface is stabilized by a distributed network of cooperative polar contacts rather than any single dominant hotspot.

### Germline-like-encoded docking geometries are distinct yet conserved across TCR lineages and HLA alleles

To assess whether the germline-mediated docking principles observed in the TCR156 system extend to other TCR-pMHC pairs, we screened a broader panel of previously characterized TCR-pMHC class I complexes following CDR3 replacement and molecular clamping (Figure S11). Representative cryo-EM 2D class averages consistent with intact TCR-pMHC-Fab assemblies were observed across multiple unrelated TCR lineages and HLA backgrounds, including HLA-A*01, HLA-A*02, HLA-A*03, HLA-A*11, HLA-A*24, and HLA-B*44. These included the A3 ^31^, DMF5 (PDB: 3QDG) ^32^, 1G4 (PDB: 2BNQ) ^33^, 21LT2-2 (PDB: 7L1D) ^34^, S8-9F3 (PDB: 8RYQ) ^35^, H27-14 (PDB: 3VXS) ^36^, and DM1 (PDB: 3DXA) ^37^ TCR systems, spanning diverse germline V-gene combinations such as TRAV21/TRBV5-1, TRAV12-2/TRBV6-4, TRAV21/TRBV6-5, TRAV12-2/TRBV9, TRAV21/TRBV9, TRAV21/TRBV7-9, and TRAV26-1/TRBV7-9. Although particle abundance and reconstruction quality varied substantially between systems, the recurrent appearance of characteristic TCR-pMHC-Fab particle architectures across unrelated receptors suggests that germline-mediated MHC engagement is broadly accessible across canonical αβ TCR lineages rather than restricted to a single TCR-HLA pair.

Among the screened systems, the DMF5 TCR formed a relatively abundant and structurally homogeneous population of complexes suitable for high-resolution reconstruction. DMF5 uses the same α-chain variable gene (TRAV12-2) as TCR156 but pairs it with a different β-chain (TRBV6-4) while recognizing the same HLA-A*02 allele. Using the clamping strategy, we determined the cryo-EM structure of the germline-like DMF5-MART1-HLA-A*02 complex at 3.1 Å resolution (**Figures 4A and S12**). The germline-like DMF5 maintained the canonical orientation of the wild-type DMF5 complex, exhibiting a nearly identical crossing angle (62.1° vs. 60.4°). Structural comparison with the wild-type DMF5 crystal structure (PDB: 3QDG) confirmed a similar docking geometry, with a coordinate RMSD of 2.60 Å upon MHC alignment (**Figure 4A**).

**Figure 4.**
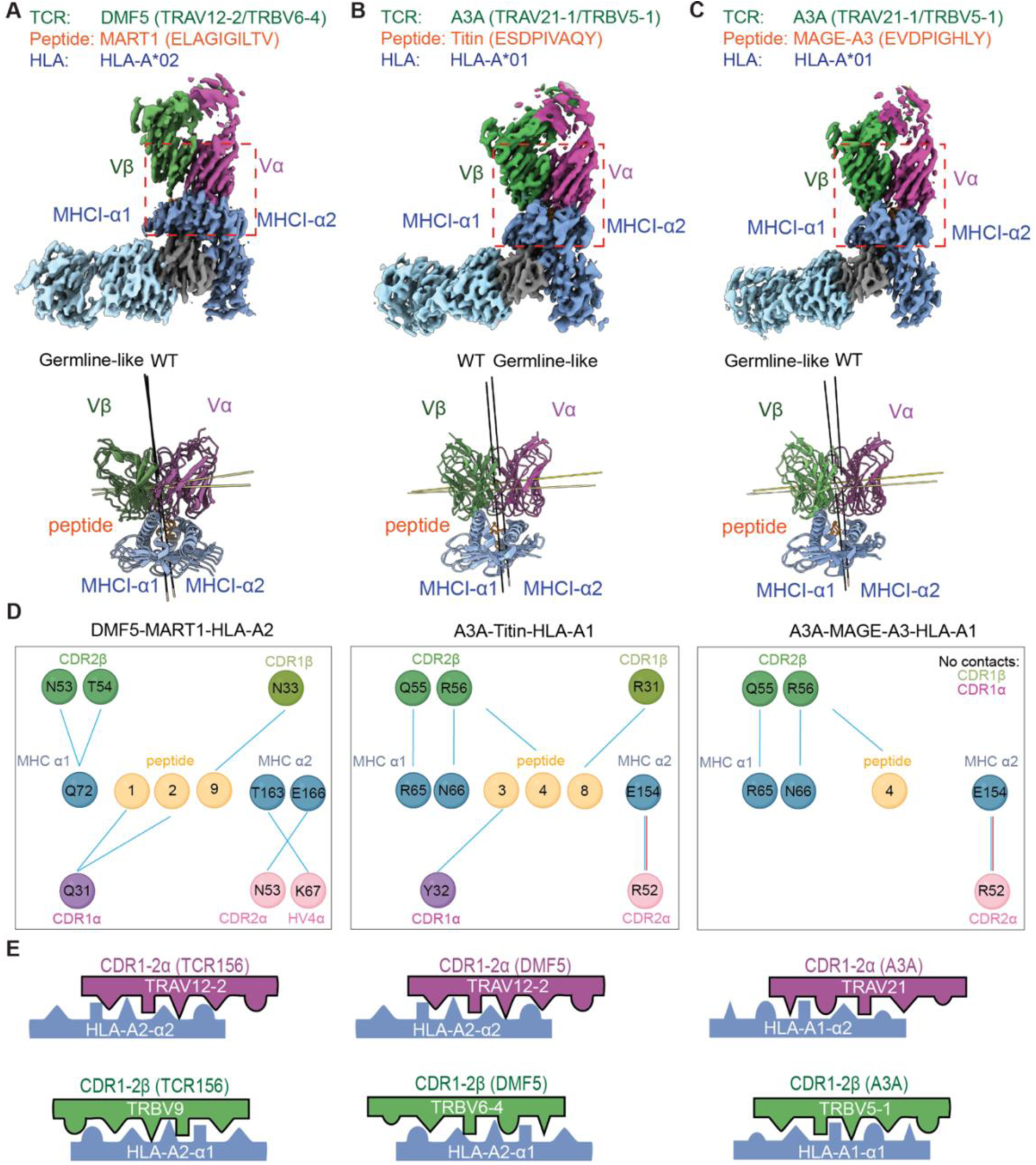
Germline-like-encoded docking geometries are distinct yet conserved across TCR lineages and HLA alleles. (A-C) Cryo-EM maps and core domain reconstructions for germline-like complexes from three TCR lineages: (A) DMF5-MART1-HLA-A*02; (B) A3A-titin-HLA-A*01; (C) A3A-MAGE-A3-HLA-A*01. For each complex, the germline-like and WT structures are shown superposed below the class averages, with TCR α (magenta), TCR β (green), peptide (orange) and MHC (blue) illustrating that the germline-like TCRs adopt the similar overall diagonal docking orientation as their WT counterparts. (D) Schematic representation of germline-mediated contacts in the DMF5 and A3A systems. The left show interactions between DMF5 and HLA-A*02 taken from the WT DMF5-MART1-HLA-A*02 crystal structure (PDB 3QDG). For consistency, DMF5 residues are renumbered in the figure to match the TCR156 TRAV12-2 numbering scheme. by applying a +1 offset to the original TCRα crystal numbering (e.g., crystal residue Q30 is renumbered as Q31). The right Two panel shows the corresponding germline contacts in the WT A3A-MAGE-A3-HLA-A*01 structure (PDB 5BRZ) and WT A3A-Titin-HLA-A*01 structure (PDB 5BS0). (E) Schematic summary of germline docking rules across lineages. For each TCR (TRAV12-2/TRBV9, TCR156; TRAV12-2/TRBV6-4, DMF5; TRAV21/TRBV5-1, A3A), the α chain (magenta) predominantly engages the MHC α2 helix, whereas the β chain (green) contacts the α1 helix, establishing a relatively conserved division of labor despite differences in Vα/Vβ usage, docking angle and peptide/HLA combination.

Although TCR156 and DMF5 both use TRAV12-2 and both recognize HLA-A*02, they adopt different overall docking angles on the MHC surface due to their distinct β-chain V-segments (TRBV9 vs. TRBV6-4) (**Figure S13**). In both receptors, TRAV12-2 residues Asn53 and Lys67 form conserved hydrogen bonds with polar residues Glu166 and Thr163 on the HLA-A*02 α2-helix, and Gln31α contacts the peptide backbone — the same contacts identified in the TCR156 series. The shared TRAV12-2 HLA-A*02 contact pattern thus acts as a conserved anchor, while the specific α/β germline combination co-defines the overall docking angle. This demonstrates that the β-chain V-segment can act as a geometric modulator that repositions the TCR heterodimer on the pMHC surface.

In contrast to DMF5, the parental A3 TCR formed only a small fraction of intact germline-like complexes, limiting direct structural analysis. We therefore utilized the affinity-enhanced A3A variant, which contains four engineered substitutions in CDR2α derived from the A3 TCR, for extended structural characterization. A3A TCR use the TRAV21/TRBV5-1 germline combination and recognizes two different peptides (MAGE-A3 and Titin) presented by HLA-A*01. Using the clamping strategy, we determined two germline-like complexes, A3A-MAGE-A3-HLA-A*01 and A3A-Titin-HLA-A*01 at 3.0 Å and 3.1 Å resolution, respectively (**Figures 4B, 4C and S14**). Comparison with the corresponding wild-type crystal structures (PDB: 5BRZ, 5BS0) the germline-like complexes exhibited highly conserved crossing angles (∼86° vs. 85° for MAGE-A3; ∼83° vs. 85° for Titin) and coordinate RMSDs of 3.8-3.9 Å upon MHC alignment ^38^.

To delineate the conserved germline-mediated contacts at atomic resolution, which we are not able to do with the lower resolution cryoEM sturctures, we utilized the available crystal structures of DMF5 (PDB: 3QDG) and A3A (PDB: 5BRZ, 5BS0) as high-resolution templates, which accurately represent the conserved interface also observed in our germline-like cryo-EM maps (**Figure 4D left**). All three structures revealed a network of hydrogen-bond interactions between germline-encoded CDR1/CDR2 loops and the MHC α-helices, consistent with the canonical TCR-MHC interface. The A3A TCR, which recognizes HLA-A*01, employs a distinct interaction network from the germline TCR interactions with HLA-A*02 (**Figure 4D, middle and right**). In this complex, Arg52α on the TRAV21 α-chain contact Glu154 on the HLA-A01 α2 helix, while Gln55β on the TRBV5-1 β-chain engage the α1 helix and Arg56β forms backbone contacts with the peptide. These allele-specific residues change remodel the local hydrogen-bond pattern without altering the basic topology of the TCR-MHC interface.

A comparative schematic of the germline interfaces (**Figure 4E**) illustrates how each TCR germline V-domain forms MHC allele-specific contacts: each TCR V-domain can form distinct MHC contacts depending on the α/β-chain pairings TRAV12-2/TRBV9 (TCR156), TRAV12-2/TRBV6-4 (DMF5), and TRAV21/TRBV5-1 (A3A). Furthermore, each TCR engages the MHC helices through a consistent division of labor, with the α-chain contacting the α2 helix and the β-chain contacting the α1 helix. These findings reinforce that MHC restriction arises from intrinsic, evolutionarily conserved features of the TCR germline, fine-tuned by specific Vα/Vβ combinations.

### Germline-encoded MHC bias is reflected in human TCR repertoire and conserved interaction motifs

To evaluate whether germline-encoded features of TCR recognition are detectable at the population level, we analyzed TCR repertoires stratified by HLA-A*02 status ^39^. Across 539 individuals, HLA-A*02-positive and -negative donors were comparably represented (**Figure 5A**). CD8⁺ clonotypes comprised a substantial fraction of the repertoire, with a large proportion residing in the naïve compartment. Because naïve CD8⁺ T cells represent a pre-selection repertoire prior to antigen-driven expansion, this subset provides a suitable framework to interrogate intrinsic, germline-level biases.

**Figure 5.**
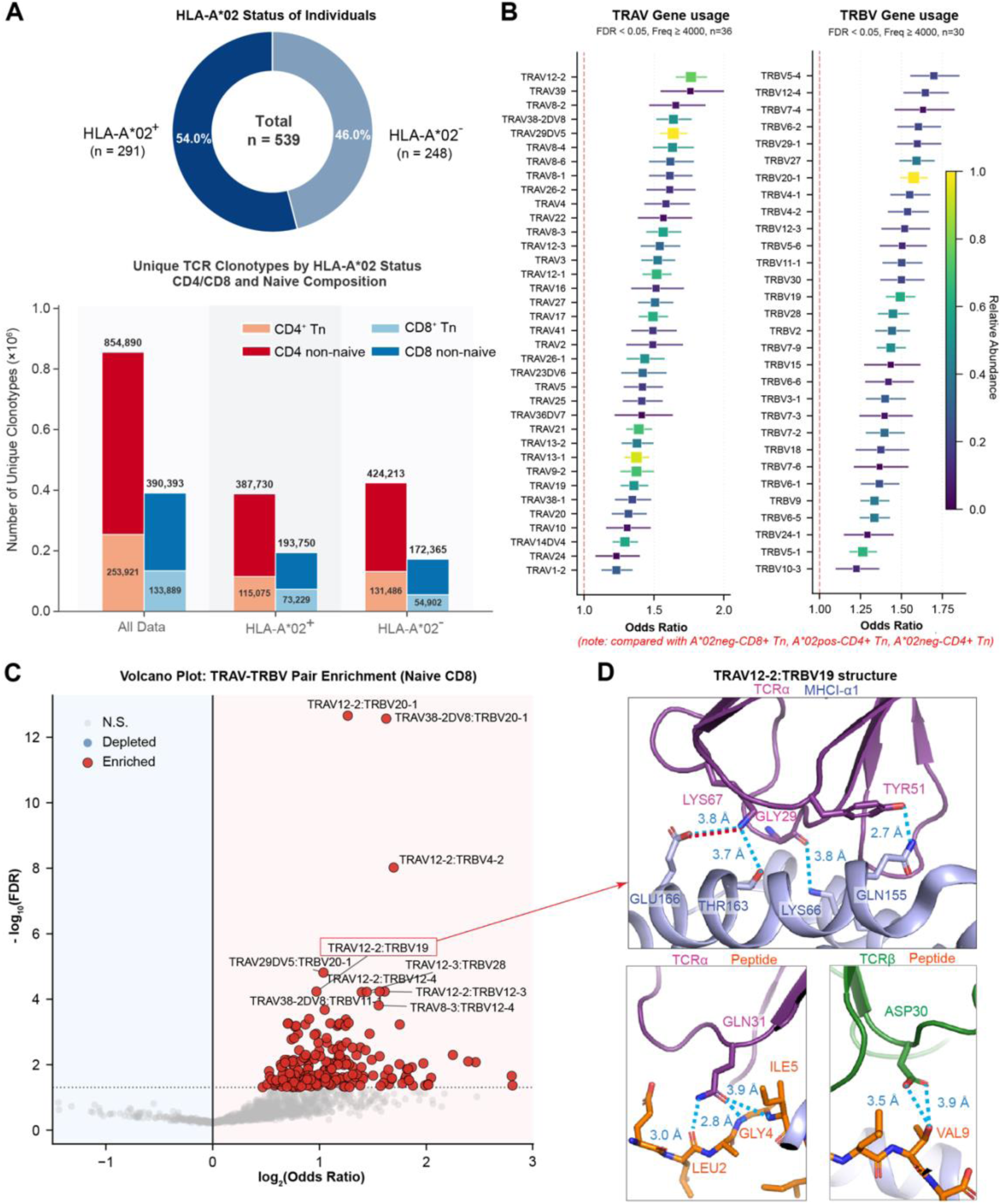
Germline-biased V gene usage and Vα-Vβ pairing patterns in HLA-A*02-restricted TCR repertoires. (A) Distribution of HLA-A*02 status across individuals (top) and the composition of unique TCR clonotypes stratified by CD4⁺/CD8⁺ and naïve (Tn) versus non-naïve subsets (bottom). Numbers above bars indicate total clonotype counts within each category. (B) Enrichment analysis of TRAV and TRBV gene usage in HLA-A*02⁺ naïve CD8⁺ T cell clonotypes relative to control groups. Points indicate odds ratios for gene usage, with horizontal bars representing confidence intervals. Only genes meeting significance (FDR < 0.05) and frequency thresholds (≥ 4,000 clonotypes) are shown. Color scale reflects proportional representation within the dataset. (C) Volcano plot of TRA-TRBV pairing enrichment in HLA-A*02⁺ naïve CD8⁺ T cell clonotypes. Each point represents a Vα-Vβ combination, plotted by log_2_(odds ratio) and −log_10_(FDR). Significantly enriched pairs (FDR < 0.05) are highlighted in red, while non-significant pairs are shown in grey. Selected pairs are highlighted for panel D. (D) Structural example linking enriched TRAV-TRBV gene pairings to germline-mediated MHC interactions. The TRAV12-2/TRBV19 TCR-pMHC complex (PDB: 5NQK) is shown, highlighting germline-encoded contacts between TCRα (purple) and the MHC helix (blue), as well as interactions involving the peptide (orange). Insets show representative hydrogen bond interactions.

At the gene level, we observed selective enrichment of specific TRAV and TRBV segments in HLA-A*02-associated repertoires. Focusing on naïve CD8⁺ clonotypes (FDR < 0.05, frequency ≥ 4,000), we identified a subset of V genes preferentially expanded in A*02-positive individuals, including TRAV12-2, TRAV39, TRAV8-2, TRAV38-2DV8, and TRAV29DV5 on the α chain, and TRBV5-4, TRBV27, TRBV12-4, TRBV29-1, and TRBV15 on the β chain (**Figure 5B**). Notably, TRAV12-2, which is used by the germline-like TCR156 and DMF5 receptors characterized structurally in this study, was among the most strongly enriched segments. A highly similar enrichment pattern was observed when the analysis was extended to the total CD8⁺ compartment (**Figure S15A**), indicating that these biases are not restricted to naïve cells but are preserved across the broader repertoire. This enrichment suggests that specific germline V gene segments are preferentially utilized in the context of particular MHC alleles.

We next asked whether such biases extend to Vα-Vβ pairing. Analysis of naïve CD8⁺ clonotypes revealed a broad set of TRAV–TRBV combinations significantly associated with HLA-A*02 (**Figures 5C and S15B**). These enriched pairs spanned a continuous range of odds ratios, indicating that V gene pairing is not random but exhibits graded preferences within the repertoire. Among the most statistically significant pairs were TRAV12-2/TRBV20-1, TRAV38-2DV8/TRBV20-1, and TRAV12-2/TRBV4-2, highlighting recurrent pairing biases within the repertoire. A highly similar pairing bias was observed when the analysis was extended to the total CD8⁺ compartment (**Figures S15C and S15D**), indicating that these pairing preferences are not restricted to naïve cells but are preserved across the broader repertoire. These included multiple TRAV12-2–containing pairs, reinforcing the gene-level enrichment and indicating that specific germline V regions contribute disproportionately to the pairing landscape.

To connect these repertoire-level observations to structural mechanisms, we examined enriched Vα-Vβ pairs with available structural information. Among these, the TRAV12-2/TRBV19 combination is represented by a solved TCR-pMHC structure (PDB: 5NQK). Structural analysis revealed that germline-encoded residues from this pairing engage the MHC helices through interaction motifs highly similar to those observed in our experimentally determined germline-like complexes, including conserved contacts between CDR1/CDR2 loops and the MHC α-helices (**Figure 5D**). These observations provide a direct structural correlate for the enrichment of specific V gene combinations in HLA-defined repertoires. These findings provide population-level support for a model in which germline-encoded features shape MHC recognition, with CDR3 diversity further contributing to peptide-dependent specificity.

## Discussion

The observation that germline-like TCRs, devoid of sequence-specific CDR3s, converge on the same MHC docking solutions as their wild-type TCR counterparts suggests a unifying model: the TCR germline encodes the blueprint for MHC restriction, upon which somatic CDR3 diversity elaborates antigen specificity. This conceptual framework is consistent with the long-standing “division of labor” model, which proposes that germline-encoded CDR1 and CDR2 loops primarily engage the MHC helices, whereas CDR3 loops focus on peptide recognition ^18,40^. Historically, the existence of germline bias has been inferred indirectly through functional and genetic analyses ^18,40^. For example, repertoire studies of HLA-A*02-restricted public TCRs, including those specific for MART1, yellow fever virus, and the SARS-CoV-2 spike protein, consistently reveal a strong TRAV12-1/12-2 bias ^41–43^. Complementary structural studies have shown that TCRs sharing related V genes engage MHC α-helices in similar orientations ^19,44–46^. However, these earlier structures were derived from fully rearranged TCRs that differed mainly in their CDR3 loops and recognized the same or related peptides, making it impossible to isolate the specific contribution of germline-encoded CDR1/CDR2 loops from peptide-dependent contacts.

Our results have practical implications for the goal of “decoding” the TCR and defining the rules by which it recognizes peptide-MHC ^47^. The advent of cancer immunotherapy and knowledge that T cells can control cancer through recognition of tumor antigens has led to initiatives to predict TCR specificity for pMHC using artificial intelligence. It is assumed that this will require the generation, and curation of very large datasets of TCR sequences with matched peptide sequences for training LLM to predict TCR specificity, which remains a distant goal ^48^. Most of these approaches focus on pairing CDR3 sequences with peptide sequences, with little consideration of germline recognition. Indeed, in most TCR-pMHC complexes, the germline components have some peptide contacts, and the CDR3s also make contacts with the MHC helices, blurring the boundaries between germline and adaptive recognition and underscoring the immense complexity of the prediction problem ^49^. Nevertheless, with the knowledge that germline-derived structural constraints exist that are independent of CDR3 and peptide, this opens the possibility of deriving a loose semblance of germline rules to serve as constraints that could simplify the peptide specificity prediction problem by CDR3 ^50^. Critically, because CDR1 and CDR2 alone reproduce the wild-type docking geometry, this constraint reflects a fixed structural property of the germline-MHC interface rather than a CDR3-dependent one, providing a more direct prior for AI-based specificity prediction than repertoire association alone. We note, however, that the TCRs characterized here were isolated as intact receptors that had already undergone thymic selection together with their native CDR3 loops; our data therefore cannot fully exclude the possibility that this germline-MHC compatibility reflects, at least in part, a signature of that selection history rather than a property of the V-gene sequence that is independent of any selection process.

It is important to consider additional factors that influence pMHC recognition by the TCR. For example, the Vα-Vβ chain pairing angles impact the subsequent CDR1 and CDR2 contact footprints on the MHC ^51^. We addressed this by comparing TCR156 and DMF5, two HLA-A*02-restricted receptors that both use TRAV12-2 yet pair with distinct β-chain variable segments (TRBV9 and TRBV6-5, respectively). These TCRs share a similar TRAV12-2 HLA-A*02 contact pattern, yet adopt different overall docking angles, indicating that the β-chain context repositions the heterodimer on the pMHC surface. Thus, TRAV12-2 provides a conserved framework for engaging the HLA-A*02 α2 helix, whereas the global geometry of TCR docking is co-defined by the specific α-β germline combination. Extending these observations, analysis of TCR156, DMF5, and A3A across distinct HLA alleles (A*02 and A*01) revealed that germline-encoded docking principles are lineage-specific in orientation. By extension, the interfacial MHC contacts for each TCR V-region are also edited by the J-regions which meet at the Vα-Vβ domain interfaces and influence the Vα-Vβ pairing angles that position the CDR loops. Across all three lineages, TCR156, DMF5, and A3A, germline residues not only contact the MHC α-helices but also form backbone-level hydrogen bonds to the peptide, providing a generic readout of peptide conformation without imposing side-chain specificity. This peptide backbone contact is likely important for stabilizing the docking footprint. Finally, as previously mentioned, CDR1 and CDR2 frequently contact peptide side chains and contribute to peptide specificity, and CDR3 contacts the MHC helices, illustrating that the respective division of labor is not absolute and is interdependent.

The observation that germline TCRs retained weak but measurable signaling activity in the absence of CDR3-peptide contacts suggests that the germline-encoded scaffold establishes the necessary alignment for signaling competence, in concert with co-receptor. While “reverse polarity” examples exist where the TCR docking footprint on the MHC is flipped from the canonical geometry ^52^, these non-canonical interactions generally do not elicit productive signaling ^53^. This may have evolutionary significance: a TCR that docks productively on MHC very weakly can be positively selected in the thymus, providing a mechanism by which MHC restriction becomes both structurally encoded and developmentally reinforced.

Consistent with these structural observations, analysis of human TCR repertoires revealed selective enrichment of specific TRAV and TRBV gene segments, as well as non-random Vα-Vβ pairing patterns, in HLA-A*02-restricted CD8⁺ naïve T cells. Notably, TRAV12-2, which is represented in the germline-like receptors characterized here, was among the most enriched Vα genes. While repertoire-level analyses cannot establish causality, these patterns are consistent with the idea that germline-encoded features contribute to shaping TCR-MHC compatibility at the population level.

The data imply that the MHC preferences of a given TCR could potentially be inferred from its V-region composition, offering a structural framework to guide computational and AI-based efforts to predict TCR-MHC compatibility directly from sequence data.

## RESOURCE AVAILABILITY

### Lead contact

Further information and requests for reagents may be directed to and will be fulfilled by K. Christopher Garcia, the lead contact.

### Materials availability

All unique reagents generated in this study are available from the lead contact with a completed Materials Transfer Agreement.

### Data and code availability

Cryo-EM maps and atomic coordinates for the SpyTag-SpyCatcher linked AS4.2-PRPF3-HLA-B*27, germline TCR156-PAP-HLA-A*02, germline TCR156-MART1-HLA-A*02, germline TCR156-NY-ESO-1-HLA-A*02, germline TCR156-poly-alanine-HLA-A*02, germline-like DMF5-MART1-HLA-A*02 germline A3A-MAGE-A3-HLA-A*01, and germline A3A-Titin-HLA-A*01 structures have been deposited in the PDB (http://www.wwpdb.org) with the accession codes 9ZET, 9ZEP, 9ZEQ, 9ZER, 9ZES, 11OE 9ZEU, and 9ZEV, and in the Electron Microscopy Data Bank (https://www.ebi.ac.uk/emdb/) with the accession codes EMD-74119, EMD-74115, EMD-74116, EMD-74117, EMD-75883, EMD-74118, EMD-74120, and EMD-74121 respectively. Custom Jupyter scripts used for TCR repertoire analysis are available on GitHub at https://github.com/Nan-Wang-2025/TCR-germline-repertoire-analysis.

## Acknowledgments

We thank B. Singal from the Stanford University Cryo-Electron Microscopy Center (cEMc), and R. Yan, J. Jung, X. Zhao, N. Spellmon from the Howard Hughes Medical Institute Janelia Research Campus Cryo-EM Facility for their assistance with data collection. Structural biology applications used in this project were compiled and configured by SBGrid. We thank members of the Garcia laboratory for support and helpful discussions; K.C.G. is an investigator with the Howard Hughes Medical Institute and is supported by NIH R01AI103867. This work was delivered as part of the MATCHMAKERS team, supported by the Cancer Grand Challenges partnership financed by CRUK (CGCATF-2023/100006), the National Cancer Institute (OT2CA297242).

## Author contributions

K.C.G conceived of and supervised the project. N.W., M. T. B., K. M. J. and K.C.G designed the experiment. N.W., M. T. B., K. M. J., X.C., C.W., X.X. and S.C.W. performed experiments for protein purifications, cryo-EM data collection and structural determination. N.W., X.C., L.D.L., R.M. and A.M. performed the T cell activation assays. W.L. and N.W. performed the TCR repertoire analysis. All authors contributed to data analysis. N.W., M. T. B., and K.C.G wrote the manuscript, and all authors edited and approved the final manuscript.

## STAR★METHODS

Detailed methods are provided in the online version of this paper and include the following:

## • KEY RESOURCES TABLE

- EXPERIMENTAL MODEL AND STUDY PARTICIPANT DETAILS
- Strains and cell culture

## • METHOD DETAILS

- TCR, pMHC, and Fab Expression and Purification
- Data collection and structure determination
- Model building and refinement
- Structural Analysis and Interaction Metrics
- T cell activation in co-culture Assays
- Surface plasmon resonance (SPR)
- TCR repertoire analysis

## • QUANTIFICATION AND STATISTICAL ANALYSIS SUPPLEMENTAL INFORMATION

Figs. S1 to S15

Tables S1 to S2

## STAR★METHODS

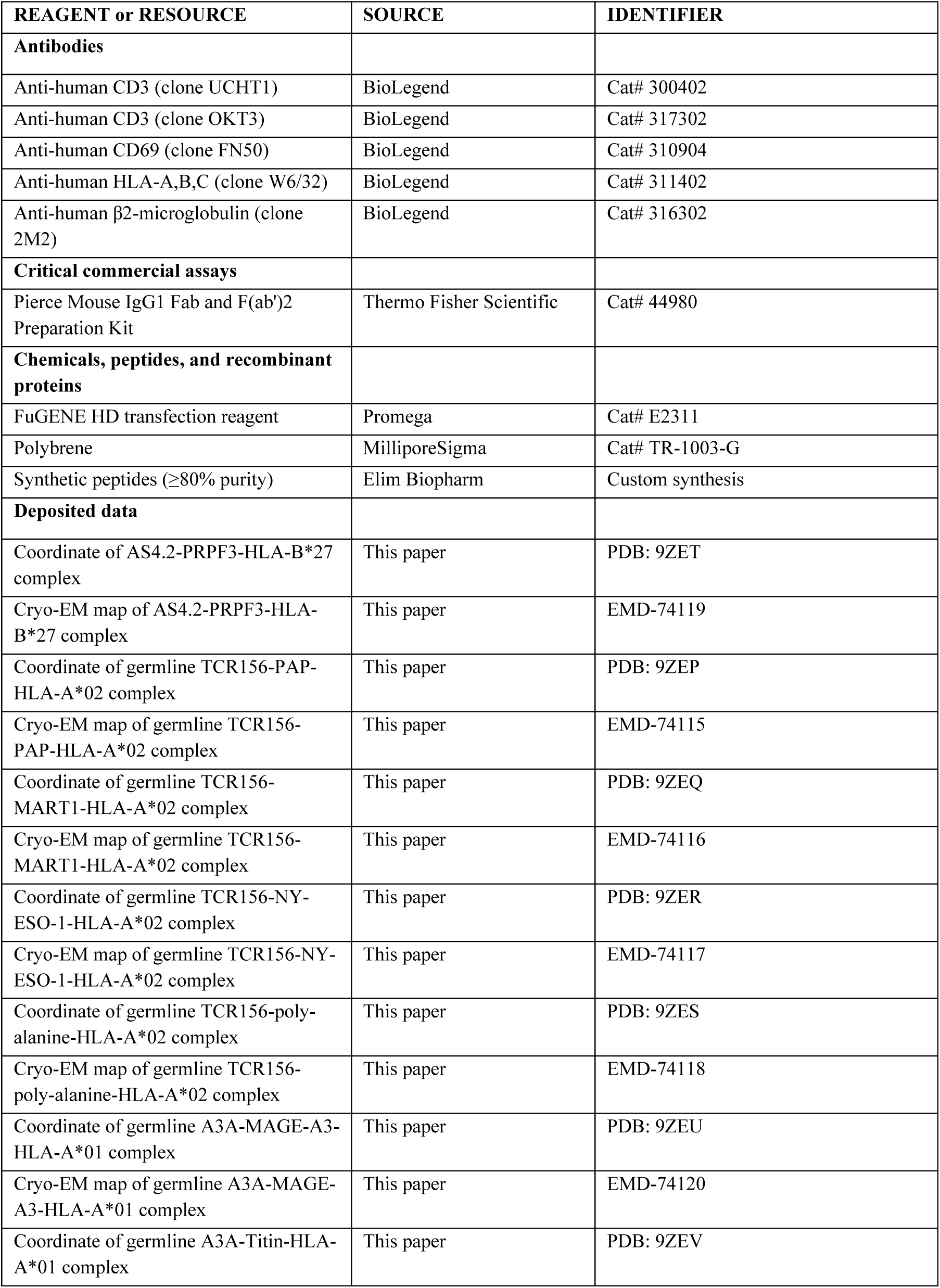
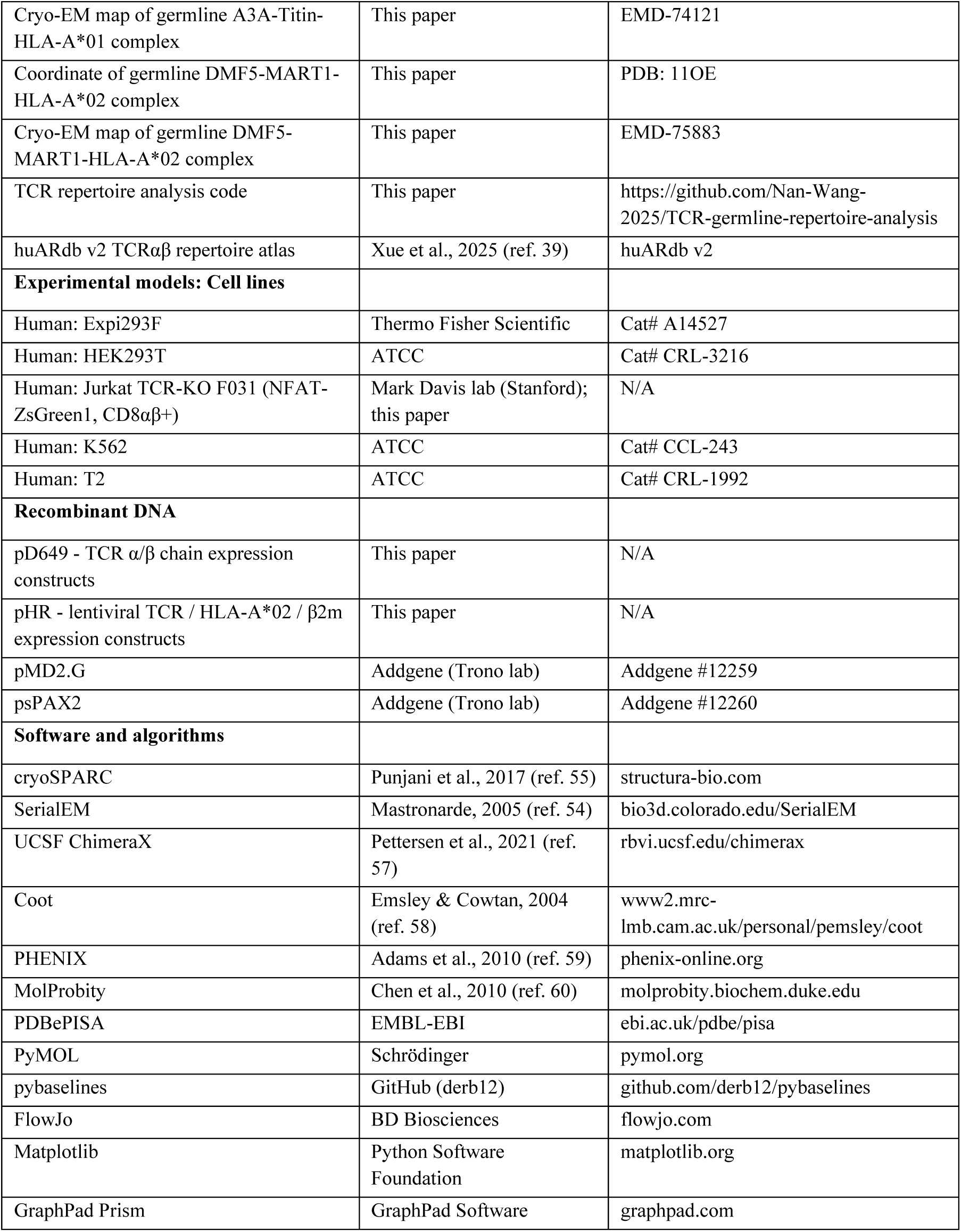
KEY RESOURCES TABLE.

## EXPERIMENTAL MODEL AND STUDY PARTICIPANT DETAILS

### Strains and cell culture

Expi293F cells were used for recombinant protein expression and were maintained in Expi293 Expression Medium with shaking. HEK293T cells were used to produce lentivirus and were cultured in DMEM supplemented with 10% FBS, penicillin, and streptomycin (P/S). F031 reporter cells were derived from a Jurkat TCR-knockout line (Mark Davis lab) engineered to constitutively express human CD8αβ and an NFAT-zsGreen1 reporter. K562 cells were transduced to express full-length HLA-A*02. T2 and K562 cells were used as antigen-presenting targets for T cell activation assays. F031, T2, and K562 cells were cultured in RPMI complete medium supplemented with 10% FBS, penicillin, streptomycin (P/S), sodium pyruvate, and L-glutamine. All cell lines were maintained at 37°C, 5% CO₂.

## METHOD DETAILS

### TCR, pMHC, and Fab Expression and Purification

The cDNAs of TCR α chains were cloned into the pD649 vector consisting of 3C protease site, a C-terminal basic helix zipper and a 6×His-tag, and β chains were cloned into the pD649 vector encoding 3C protease site, a C-terminal acidic helix zipper and a 6×His-tag. TCRs α and β chains were co-transfected and co-expressed in the Expi293F system. Modifications included the addition of SpyCatcher to the N-terminus of the TCR β chain linked by flexible Gly/Ser linkers of 13-amino acid lengths, respectively.

Single Chain pMHC were cloned into the pD649 vector in the format peptide- (G₃S)₃-β₂m-(G₄S)₄-HLA-α-10×His. For Spy-tag HLA, a spy-tag was adding after (G_3_S)_3_ linker, and an additional 8-residue Gly/Ser linker was added before β_2_M. This strategy was used to generate all pMHC constructs used in this study.

Following a four-day post-transfection period, the cell culture was centrifuged at 2,000 g for 10 minutes to collect the supernatant, which was then mixed with an equal volume of PBS pH7.2 (Gibco) and 2 mL of Ni-NTA slurry (Qiagen) per 50 mL of supernatant. This mixture was rotated overnight at 4°C. Then, the solution was passed through a column to collect the Ni-NTA and bounded protein. The resin was washed with 30 column volumes of PBS pH 7.2 containing 20 mM imidazole and eluted with 10 column volumes of PBS pH 7.2 containing 200 mM imidazole. After concentration through a 30-kDa filter (Millipore). Zippers from TCRs were cleaved using a 3C protease (1:50 w/w ratio) overnight at 4°C. Final purification was performed via size-exclusion chromatography on a Superdex 200 column (GE Healthcare) equilibrated with PBS pH 7.2. β2M Fab was obtained by digesting anti-human β2M antibody (Clone 2M2, Biolegend) using the Pierce Mouse IgG1 Fab and Preparation Kit (Thermo Fisher).

Peak fractions containing purified TCRs and pMHCs were collected and concentrated separately. Protein concentrations were determined using a Nanodrop spectrophotometer (Thermo Fisher Scientific). Purified TCR, pMHC, and Fab components were then mixed in a 1:1:1.2 molar ratio and incubated on ice for one hour to promote complex formation. The assembled complexes were subjected to a final purification step by size-exclusion chromatography on a Superdex 200 Increase column (GE Healthcare) equilibrated with PBS (pH 7.2). Peak fractions corresponding to the ternary complex were pooled and concentrated to approximately 3-10 mg/mL for cryo-EM grid preparation and analysis.

### Data collection and structure determination

To prepare the cryo-EM specimens, 3.0 μL of each protein complex was applied to a glow-discharged grid (Quantifoil Au R1.2/1.3, 200 mesh) and excess sample was blotted by a filter paper for 1.0 s before plunge-freezing in liquid ethane cooled by liquid nitrogen with Vitrobot (Mark IV, Thermo Fisher Scientific) at 8°C and 100% humidity. Cryo-EM movies were collected using an FEI Titan Krios operated at 300 kV equipped with Gatan K3 camera.

Micrographs of SpyTag-SpyCatcher linked AS4.2-PRPF3-HLA-B*27, germline TCR156-PAP-HLA-A*02, germline TCR156-MART1-HLA-A*02, germline TCR156-NY-ESO-1-HLA-A*02, germline TCR156-poly-alanine-HLA-A*02, germline DMF5-MART1-HLA-A*02, germline A3A-MAGE-A3-HLA-A*01, germline A3A-Titin-HLA-A*01 were recorded using SerialEM ^54^ in the super-resolution mode with a pixel size of 0.426, 0.4195, 0.4285, 0.4135, 0.4285, 0.426, 0.4285, and 0.4285Å. Patch motion correction was performed with binning to the physical pixel sizes of 0.852, 0.839, 0.857, 0.827, 0.857, 0.852, 0.857, and 0.857Å. Each stack of 50 frames was exposed with total dose 60 e-/Å2 at the specimen, and the defocus range between -1.0 and -2.0 μm. All collected movies were processed and assessed with cryoSPARC Live ^55^.

For the AS4.2-PRPF3-HLA-B*27 dataset, 5,491 non-tilt micrographs and 2,115 tilt micrographs with were collected and used for the downstream processing. For non-tilt data, 7,303,614 particles were initially picked and extracted in a 280-pixel box (binned). After three rounds of 2D classifications and seed-facilitated guided multi-reference 3D classification ^56^, 1,288,857 particles re-extract with the 280/200 binned box are selected followed by refinement generating a 2.8 Å map. For tilt data, 384,170 particles generated a 4 Å map using the same box size. Non-tilt and tilt data are combined, after 3D classification with resolution gradient maps and Local refinement with a core domain mask push the density resolution to 2.8 Å.

The germline TCR156-HLA-A*02 peptide series (PAP, MART1, NY-ESO-1 and poly-alanine) was processed using an analogous workflow (**Figures S7 and S8**). For GL156-PAP-A2, GL156-MART1-A2, GL156-NYESO-A2 and GL156-polyA-A2, 9,893, 5,535, 5,679 and 4,803 micrographs were collected, respectively. Particles were initially identified by template or blob picking and extracted in binned boxes, yielding 7,368,688 (PAP), 10,621,216 (MART1), 5,387,521 (NYESO) and 5,205,107 (polyA) particles. After two rounds of 2D classification, ab initio reconstruction and three rounds of guided multi-reference 3D classification, 1,606,278 (PAP), 355,911 (MART1), 578,403 (NYESO) and 373,586 (polyA) particles were retained for further refinement. These particles were re-extracted into 320 px to 240 px boxes and subjected to resolution-gradient 3D classification to remove low-resolution and junk classes. The final selected particle sets (1,217,845 for PAP, 224,288 for MART1, 293,406 for NYESO and 274,702 for polyA) were refined by global and local CTF refinement, non-uniform refinement and local refinement with a core-domain mask, followed by map sharpening, resulting in overall resolutions of 2.9 Å (PAP), 3.1 Å (MART1), 3.0 Å (NYESO) and 3.2 Å (polyA).

The germline DMF5-MART1-HLA-A*02 dataset (GLDMF5-MART1-A2) was collected and processed using the same cryoSPARC workflow as the TCR156 peptide series (**Figure S12**). Briefly, 5,162 micrographs were acquired, and blob-picked particles were extracted in binned 320-pixel boxes. After two rounds of 2D classification to remove obvious contaminants and ice, the remaining particles were subjected to ab initio reconstruction followed by multiple rounds of guided multi-reference 3D classification. A single well-resolved class was selected and re-extracted in larger boxes at the physical pixel size. Global and local CTF refinement and non-uniform refinement produced a final reconstruction at 3.10 Å resolution.

For the HLA-A*01 complexes, germline A3A-MAGE-A3-HLA-A*01 (GLA3A-MAGE-A1) and germline A3A-titin-HLA-A*01 (GLA3A-Titin-A1) were each collected as paired non-tilted and tilted datasets (**Figure S14**). Movies were imported into cryoSPARC, motion-corrected and CTF-estimated as described above. For GLA3A-MAGE-A1, particles were picked separately from the non-tilted and tilted micrographs, combined, and subjected to iterative 2D classification to discard junk and broken particles. Good particles were used for ab initio 3D reconstruction and heterogeneous refinement, yielding a dominant class corresponding to the intact GLA3A-MAGE-A1 complex. These particles were re-extracted in unbinned boxes and refined by non-uniform refinement, followed by local CTF refinement and local refinement with a core-domain mask, resulting in a final map at 3.0 Å resolution. The GLA3A-titin-A1 dataset was processed analogously. Non-tilted and tilted movies were processed in parallel, and good particles from both were combined. Using the low-pass-filtered (12 Å) GLA3A-MAGE-A1 map as the initial reference for non-uniform refinement yielded a 3.1 Å reconstruction.

The resolution of all datasets was estimated using the gold-standard Fourier shell correlation (FSC) 0.143 criterion, and the angular distributions of the particles used for the final reconstructions showed some levels of anisotropy, indicative of the sample orientations during imaging.

### Model building and refinement

The atomic coordinates of all TCR-pMHC complexes were generated by homology modeling using high-resolution crystal structures as templates (Supplementary Tables 1 and 2). For AS4.2-PRPF3-HLA-B*27, the crystal structure of the same complex (PDB 7N2N) was used as the starting model. For the germline TCR156-HLA-A*02 series, we used the WT TCR156-PAP-HLA-A*02 structure (PDB 9NMU) as the base template, in combination with peptide/MHC coordinates from related structures where appropriate (MART1 from PDB 3QDG and NY-ESO-1 from PDB 9C3E). For germline DMF5-MART1-HLA-A*02, the DMF5-HLA-A*02 crystal structure (PDB 3QDG) was used as the initial model, and for the germline HLA-A*01 complexes we used the A3A-MAGE-A3-HLA-A*01 and A3A-titin-HLA-A*01 crystal structures (PDB 5BRZ and 5BS0, respectively).

Each model was individually fitted into the corresponding cryo-EM density map by rigid-body docking in UCSF ChimeraX ^57^, then manually adjusted and rebuilt in COOT ^58^, followed by real-space refinement in PHENIX ^59^. Validation of the models included checking MolProbity scores and Ramachandran plot statistics ^60^, with scores calculated according to established methods (Supplementary Tables 1 and 2).

### Structural Analysis and Interaction Metrics

To quantitatively compare the docking geometry and interface properties of all TCR-pMHC structures, we performed multi-scale structural analyses. MHC residues are numbered according to the standard HLA-A*02 (or HLA-A*01) mature polypeptide sequence; although single-chain trimer (SCT) constructs were used for structural studies. To assess the relative displacement of the TCR on the MHC platform, the MHC α1-α2 helix were used as a reference for superimposition, followed by a Cα root-mean-square deviation (RMSD) calculation for the TCR variable domains (Vα and Vβ). Crossing angles were determined by measuring the orientation of the TCR variable domain pseudosymmetry axis relative to the MHC peptide-binding groove.

To evaluate the structural and biophysical congruence between the wild-type (WT) and germline-like complexes, metrics were normalized and visualized using a multi-dimensional radar plot. For the radar plot, the intrinsic structural match of the Vα and Vβ domains was determined by calculating the Cα RMSD after optimal structural alignment (align) and converted to a similarity score using the formula: 1-(RMSD/5.0 Å). For docking geometry, the similarity score was calculated as 1 - | (θ_WT_ – θ_Germline_)/θ_WT_ |. The total BSA at the TCR-pMHC interface was calculated using the measure buried area command in ChimeraX with a probe radius of 1.4 Å. Inter-molecular contacts were identified using mainly PDBePISA, but also inspected using PyMol contacts feature and with manual inspection of the structures, using a distance cutoff of 3.6 Å between atoms of the TCR and the pMHC, unless indicated otherwise in the figure legend. To quantify the contribution of germline-encoded “interaction motifs” to interface stability, we defined core motifs as TCR residues in CDR1/2 loops that form recurrent contacts with the MHC helices. The Interaction Motif Coverage Index was calculated as the ratio of the BSA contributed by these specific residues in the germline-like complex relative to the WT complex. All structural measurements and visualizations were performed in UCSF ChimeraX.

### T cell activation in co-culture Assays

Full-length TCR α- and β-chains were cloned into separate pHR lentiviral vectors. Lentiviral particles were produced by co-transfecting HEK293T cells with the TCR expression plasmid, pMD2.G, and pSPAX2 using FuGENE HD (Promega) in DMEM supplemented with 10% FBS. Viral supernatants were collected 48 h post-transfection, clarified by centrifugation, and used immediately or stored at -80 °C.

F031 reporter cells were generated from the Jurkat TCR-knockout line (provided by the Mark Davis laboratory) and engineered to constitutively express human CD8αβ and an NFAT-zsgreen1 reporter. For transduction, 1.5×10⁶ F031 cells were spinfected with 1 mL lentiviral supernatant in the presence of 8 µg/mL polybrene (MilliporeSigma). Cells were cultured in complete RPMI and expanded for 5-7 days. TCR surface expression was confirmed by flow cytometry using anti-CD3 antibodies (UCHT1, BioLegend).

Full-length HLA-A*02 variants and β2-microglobulin (β2m) were cloned into the pHR vector and packaged using the same lentiviral production protocol. Viral supernatants were used to transduce 1.5×10⁶ K562 cells by spinfection. Following 5-7 days of expansion, surface MHC expression was verified by staining with anti-HLA-A, B, C (W6/32, BioLegend).

For peptide presentation, T2 cells or HLA-A*02-transduced K562 cells were incubated with peptides at the indicated concentrations for 2-3 h at 37 °C, 5% CO₂. Peptides (≥80% purity; Elim Biopharm) were dissolved at 10 mM in DMSO and diluted in RPMI to the desired working concentration. After peptide loading, cells were washed once with RPMI and co-cultured with TCR-expressing F031 cells at a 1:1 effector:target ratio for 14-18 h.

T cell activation was assessed by staining for CD69 (FN50, BioLegend) and CD3 (OKT3 or UCHT1, BioLegend), along with detection of NFAT-zsgreen1 fluorescence (FITC channel). Samples were analyzed on a CytoFLEX flow cytometer (Beckman Coulter), and data were processed using FlowJo (BD).

### Surface plasmon resonance (SPR)

The peptide-MHC-β2M single chain trimer, tagged with BirA, is initially purified using a nickel-NTA column. Subsequently, the proteins are biotinylated overnight and further purified by size-exclusion chromatography using a Superdex 200 column on an ÄKTA pure FPLC system (GE Healthcare). TCR affinity to the specific pMHC was measured by SPR on a Biacore T100 (GE Healthcare). The biotinylated single chaing trimer was immobilized on streptavidin chip (GE Healthcare) until 100-200 RU. The TCR protein was treated with 3C protease to remove the basic/acid zipper. The pMHC protein was immobilized until a 150-250 RU increase, and serial dilutions of TCR protein were flowed through the flow cell at 25°C. Baseline drift was corrected using the mixture model function of the pybaseline package (https://github.com/derb12/pybaselines).

Stable binding RU were plotted as a function of TCR concentration, and a 4-parameter logistic model was fitted to the data.

### TCR repertoire analysis

TCR repertoire analyses were performed from huARdb v2 TCRαβ repertoire atlas containing paired TCR annotations, cell subtype annotations, individual identifiers, and HLA genotypes ^39^. HLA-A*02 status was assigned per individual according to whether either A_1 or A_2 contained an A*02 allele. For naïve-compartment analyses, clonotypes annotated as CD8^+^ Tn or CD4^+^ Tn were retained.

For TRAV and TRBV enrichment analyses, clonotype counts were tabulated across four groups: A02-positive CD8, A02-positive non-CD8, A02-negative CD8, and A02-negative non-CD8, or, in naïve-restricted analyses, A02-positive CD8^+^ Tn, A02-positive CD4^+^ Tn, A02-negative CD8^+^ Tn, and A02-negative CD4^+^ Tn. For each gene, enrichment in the A*02-positive CD8 compartment was assessed using 2 × 2 contingency tables and Fisher’s exact test, followed by Benjamini-Hochberg correction. Genes with FDR < 0.05 and total frequency ≥ 4,000 were considered significant and displayed.

For Vα-Vβ pairing analyses, TRAV-TRBV combinations were defined by concatenating the annotated TRAV and TRBV genes for each clonotype. Pair counts were analyzed using the same 2 × 2 framework and statistical procedure. For visualization, only pairs with total count ≥ 50 were retained. Odds-ratio plots, volcano plots, and lollipop plots were generated in Python using Matplotlib.

## QUANTIFICATION AND STATISTICAL ANALYSIS

Data were analyzed using Python (SciPy) and GraphPad Prism. Gene and gene-pair enrichment in TCR repertoires was assessed by Fisher’s exact test with Benjamini-Hochberg correction; differences were considered statistically significant at FDR < 0.05. SPR binding curves were fit with a 4-parameter logistic model in Prism.

