## Supplementary Materials for "MHC restriction is structurally encoded by the T cell receptor germline"

#### **Summary**

MHC restriction — the requirement for $\alpha\beta$ T cell receptors (TCRs) to recognize Major Histocompatibility Complex (MHC) molecules when presenting peptide antigens — is a pillar of adaptive immunity. Here, we provide direct structural evidence for TCR germline-encoded recognition of MHC molecules irrespective of antigenic peptide. We engineered "germline-like" TCRs to eliminate CDR3 specificity for peptide, and a molecular clamping strategy to trap ultra-low-affinity complexes with peptide-MHC for cryo-EM. We find that germline-like TCR/pMHC docking modes are identical to wild-type T cell receptors with intact CDR3 loops, across distinct peptide antigens and MHC alleles. Analysis of human TCR repertoires reveals enrichment of specific V genes and $V\alpha$-$V\beta$ pairing patterns associated with HLA-A\*02 contexts. Thus, the TCR germline encodes the blueprint for MHC restriction, providing a sequence-based framework that could inform computational and AI-driven prediction of TCR specificity.

#### **The PDF file includes:**

Figures. S1 to S15
Tables S1 to S2

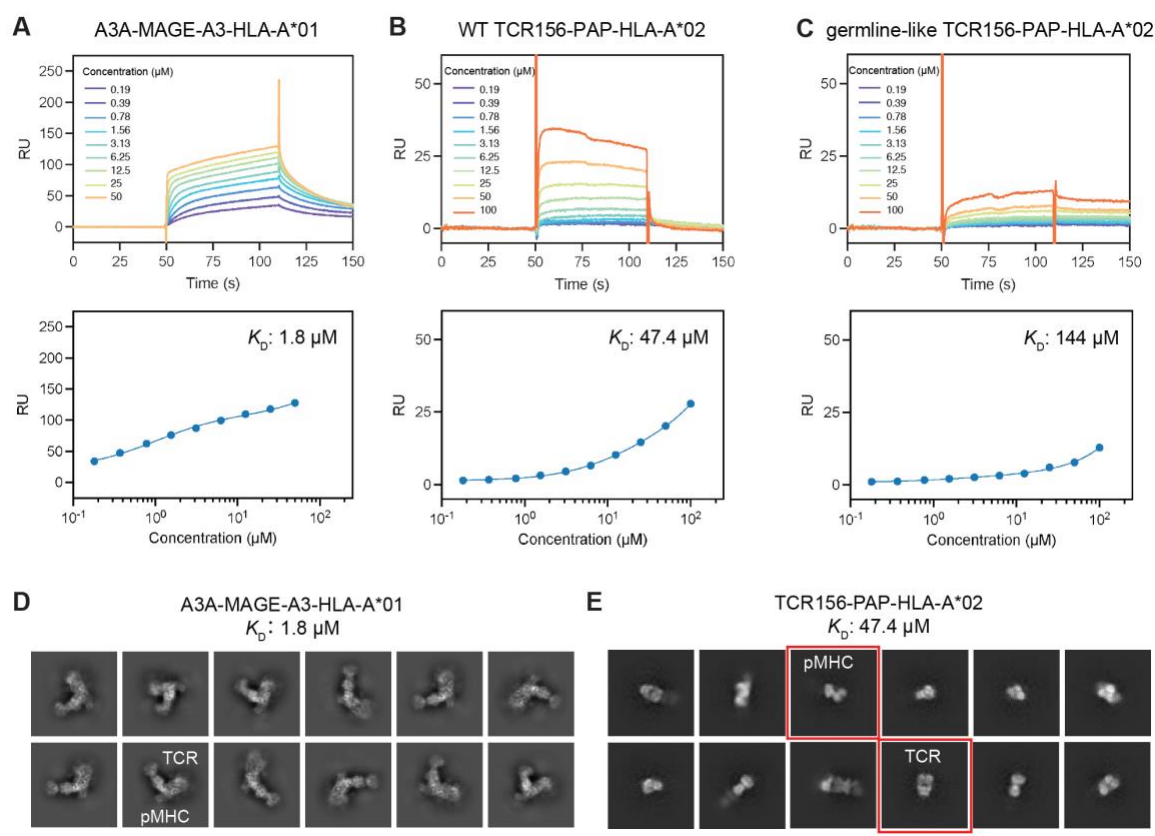

**Figure S1. SPR analysis and cryo-EM assessment of TCR-pMHC complex formation.**

(A-C) Surface Plasmon Resonance (SPR) analysis of TCR interactions. (A) Represents an example of “higher affinity” TCR-pMHC complex as measured for WT TCR A3A and MAGE-A3-HLA-A\*01 at 1.9 μM by SPR. (B) Shows the binding affinity of WT TCR156-PAP-HLA-A\*02 at 47.4 μM. (C) Illustrates the binding affinity of germline-like TCR156-PAP-HLA-A\*02 at 144 μM.

(D-E) Representative 2D class averages from cryo-EM data collection of the corresponding complexes. (D) The “higher-affinity” A3A-MAGE-A3-HLA-A\*01 complex yielded well-defined, intact TCR-pMHC particles, consistent with its tighter binding affinity. (E) In contrast, the low-affinity TCR156-PAP-HLA-A\*02 complex showed extensive dissociation, with separate populations of unbound pMHC and TCR visible in the micrographs and marked with red squares.

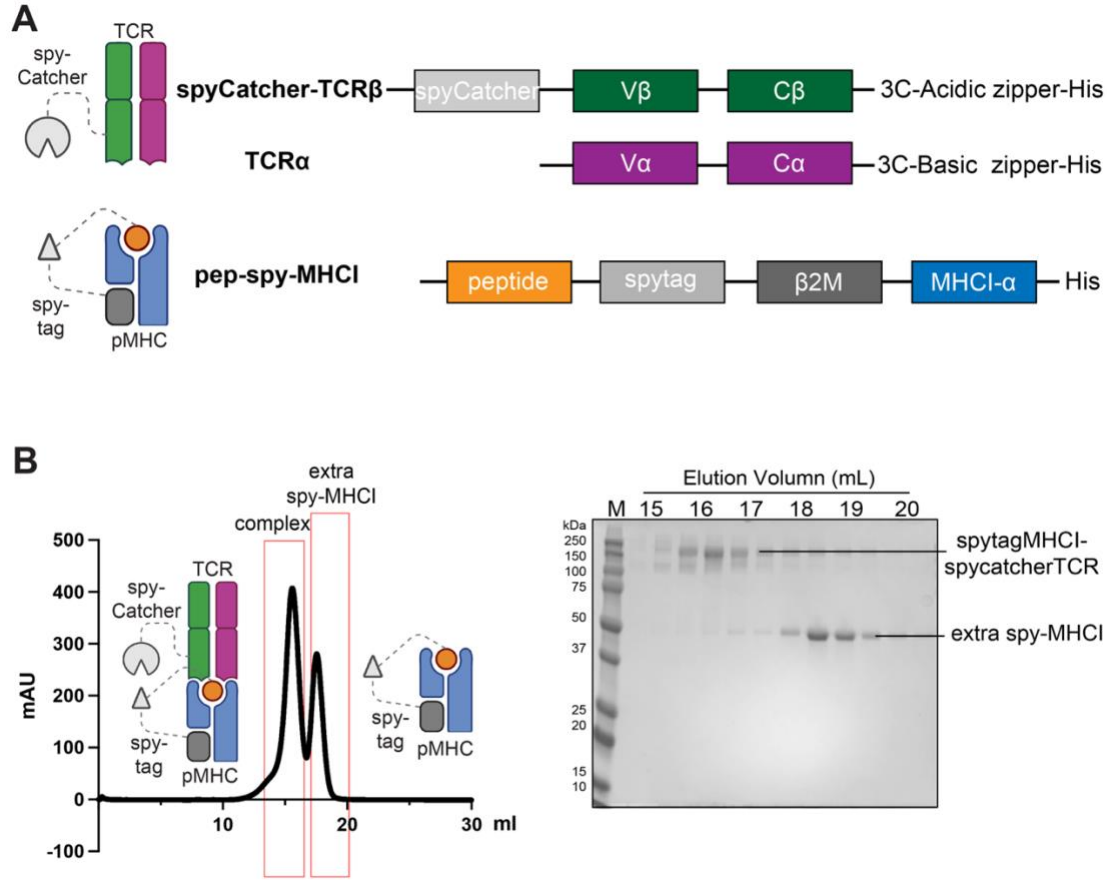

**Figure S2. Spy-tag/spyCatcher clamping strategy for TCR-pMHCI.**

(A) Schematic of the constructs used for clamping. SpyCatcher is fused to the N terminus of TCRβ, which is co-expressed with the matching TCRα chain bearing complementary acidic/basic zipper-3C-His tags. The single-chain pMHCI contains the peptide, followed by SpyTag, β2m and the HLA-A\*02 α chain with a C-terminal His tag. All constructs include signal peptides and were expressed in Expi293F cells.

(B) Size-exclusion chromatography (SEC) trace of a representative clamped complex formed between SpyCatcher-TCR156 and SpyTag-PAP-HLA-A\*02. SDS-PAGE of SEC fractions (right) show a single dominant band for the TCR-pMHCI complex, confirming efficient and stoichiometric complex formation.

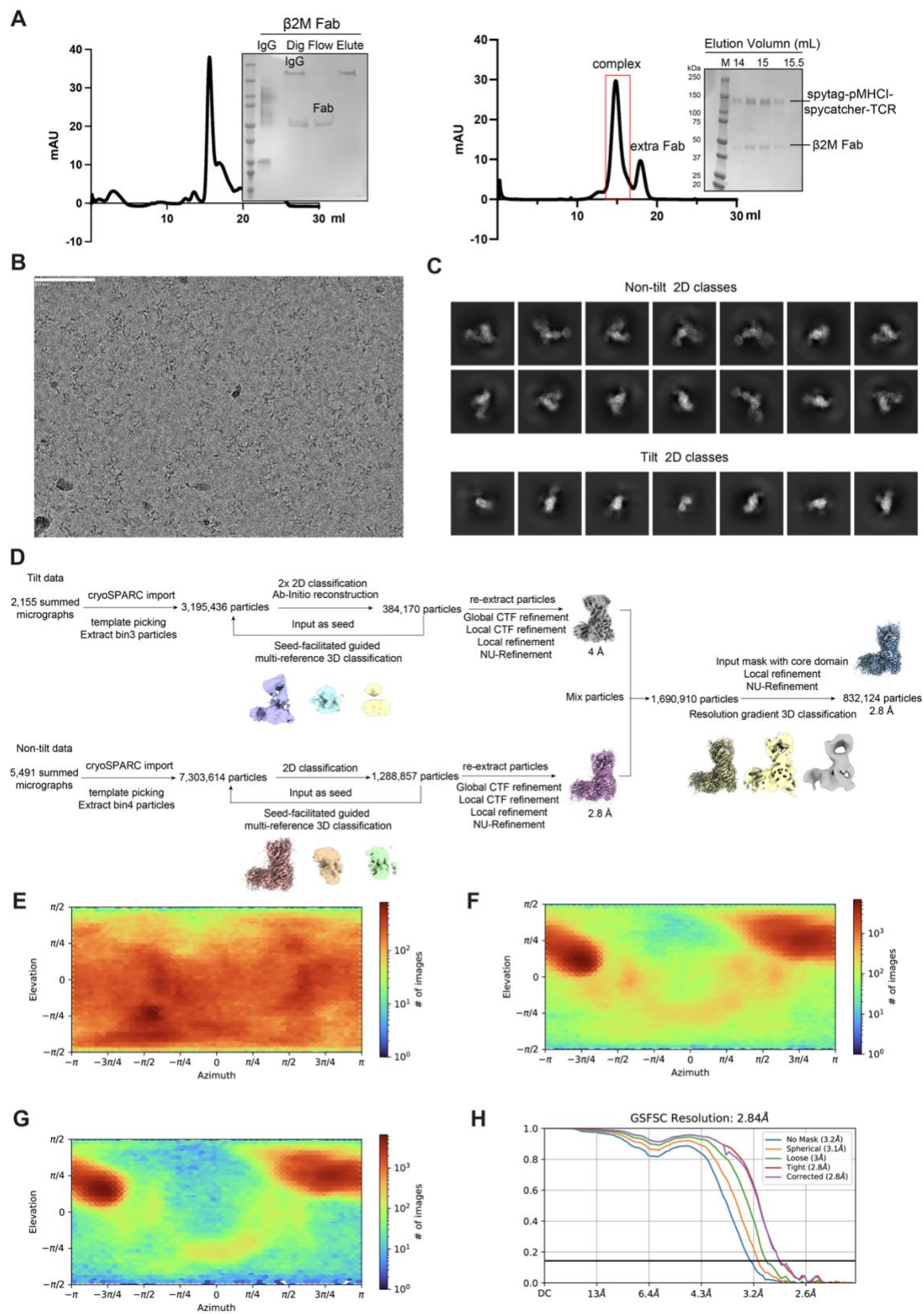

**Figure S3. Purification and cryo-EM data processing for AS4.2-PRPF3-HLA-B27\*05 complex with clamping strategy.**

(A) Representative size exclusion chromatogram of β2M Fab and AS4.2-PRPF3-HLA-B\*27 complex with clamping strategy is shown. Peak fractions were pooled and examined by Coomassie blue staining of SDS-PAGE.

(B) Representative micrograph of the AS4.2-PRPF3-HLA-B\*27 complex with clamping strategy. Scale bar = 80 nm.

(C) Representative 2D class averages showing various orientations of the AS4.2-PRPF3-HLA-B\*27 complex, including both tilt and non-tilt data.

(D) Data processing workflow for the AS4.2-PRPF3-HLA-B\*27 complex using cryoSPARC.

(E-G) Angular distribution of particles used for the final reconstructions of the AS4.2-PRPF3-HLA-B\*27 complex, with (E) showing tilt data, (F) showing non-tilt data, and (G) displaying combined tilt and non-tilt data.

(H) FSC curve for the AS4.2-PRPF3-HLA-B\*27 complex indicating a final resolution of 2.8 Å at the FSC 0.143 criterion.

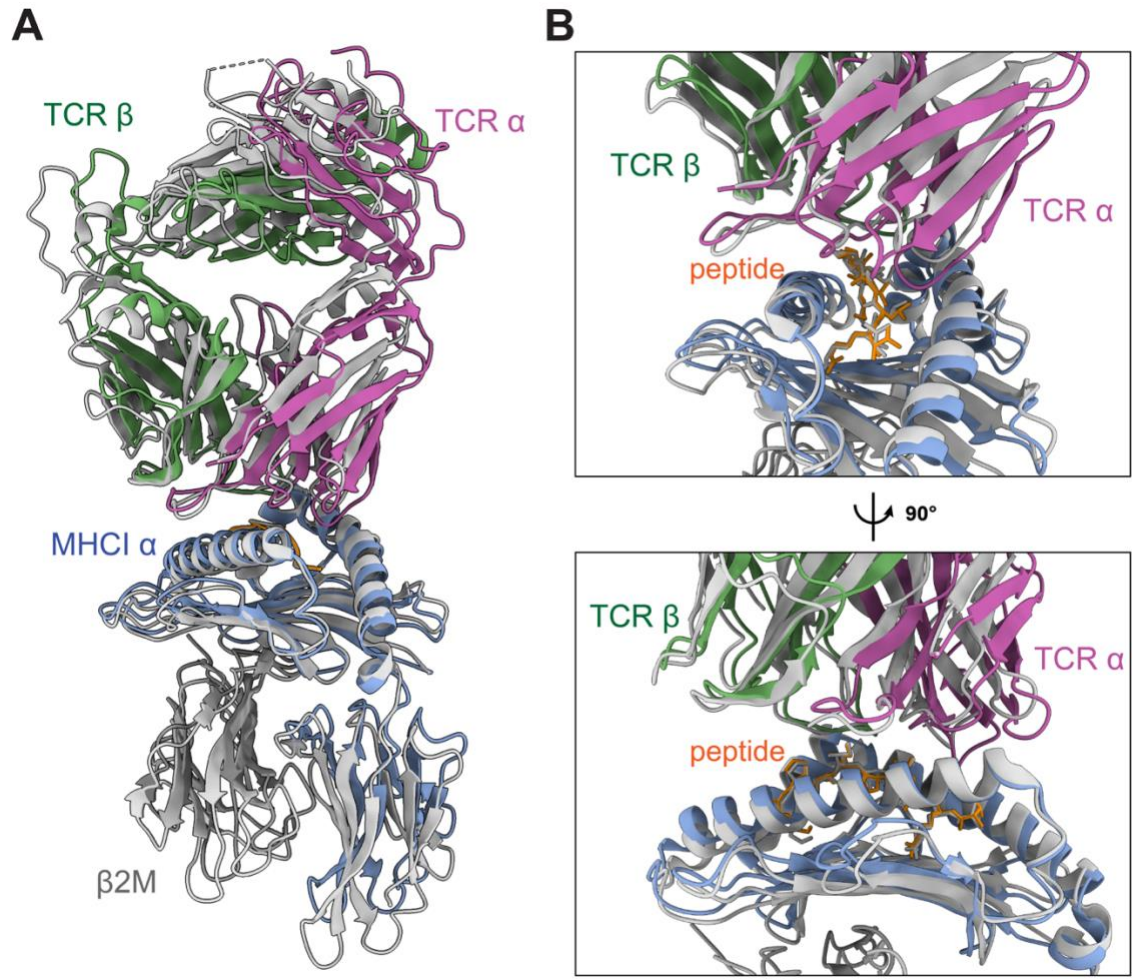

**Figure S4. Structural comparison of TCR-pMHC complexes stabilized by clamping with corresponding crystal structures.**

(A) Overall comparison of the cryo-EM structure of the AS4.2-PRPF3-HLA-B\*27 complex (colored; TCRα in magenta, TCRβ in green, peptide in orange, HLA-A\*02 $\alpha$ chain in blue), stabilized using the clamping strategy with β2M-specific Fab, and the corresponding crystal structure (colored as gray; PDB: 7N2N). Structures were aligned on the HLA-B\*27 α1-α2 platform. The MHC-aligned TCR variable domain Cα RMSD is 2.22 Å (229 atom pairs), and the crossing angles are 49.47° (cryo-EM) and 44.69° (crystal), indicating preservation of the overall docking geometry.

(B) Close-up views of the TCR-peptide-MHC interface, shown from two orthogonal orientations (top and bottom).

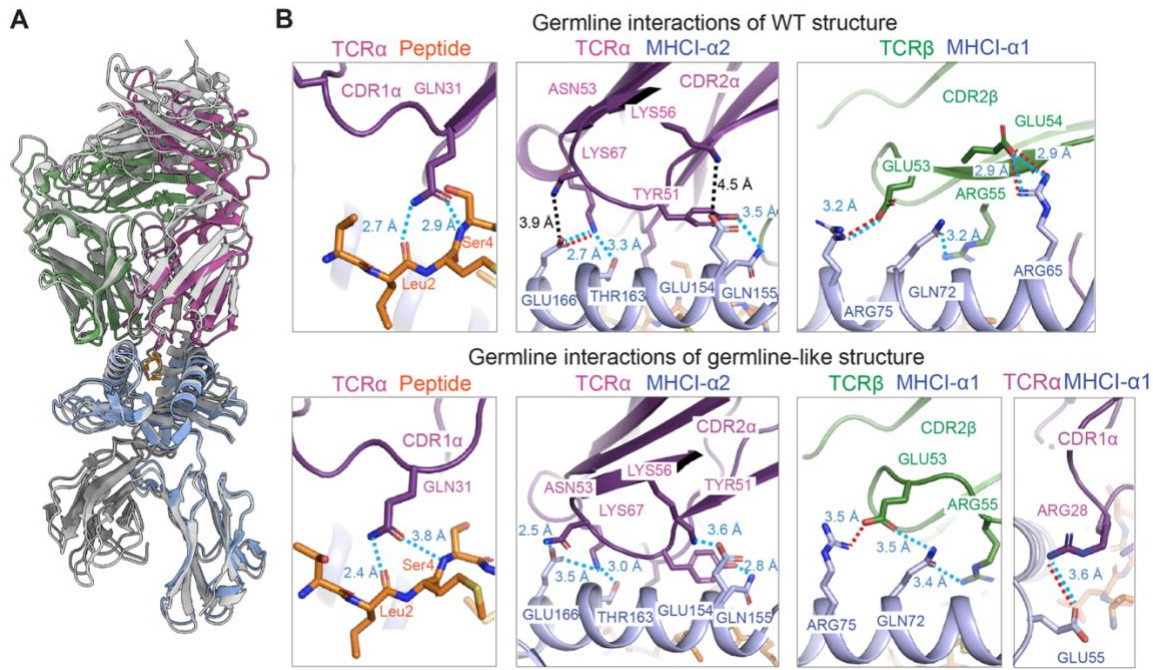

**Figure S5. Structural comparison of WT and germline TCR156-PAP-HLA-A\*02.**

(A) Superposition of the WT TCR156-PAP-HLA-A\*02 crystal structure (colored as gray; PDB: 9NMU) with the germline-like complex (colored TCRα in magenta, TCRβ in green, peptide in orange, HLA-A\*02 α chain in blue) shows that both adopt a nearly identical docking geometry. Structural alignment of the MHC α1- α2 helices reveals a coordinate RMSD of 2.20 Å for the TCR variable domains, while direct alignment of the Vα and Vβ frameworks shows an intrinsic structural match of 1.18 Å.

(B) Close-up views of germline-mediated contacts mapped onto the WT and germline-like structures, highlighting conserved CDR1/CDR2 interactions with the MHC α1/α2 helices and a backbone-level interaction between CDR1α Gln31 and the PAP peptide.

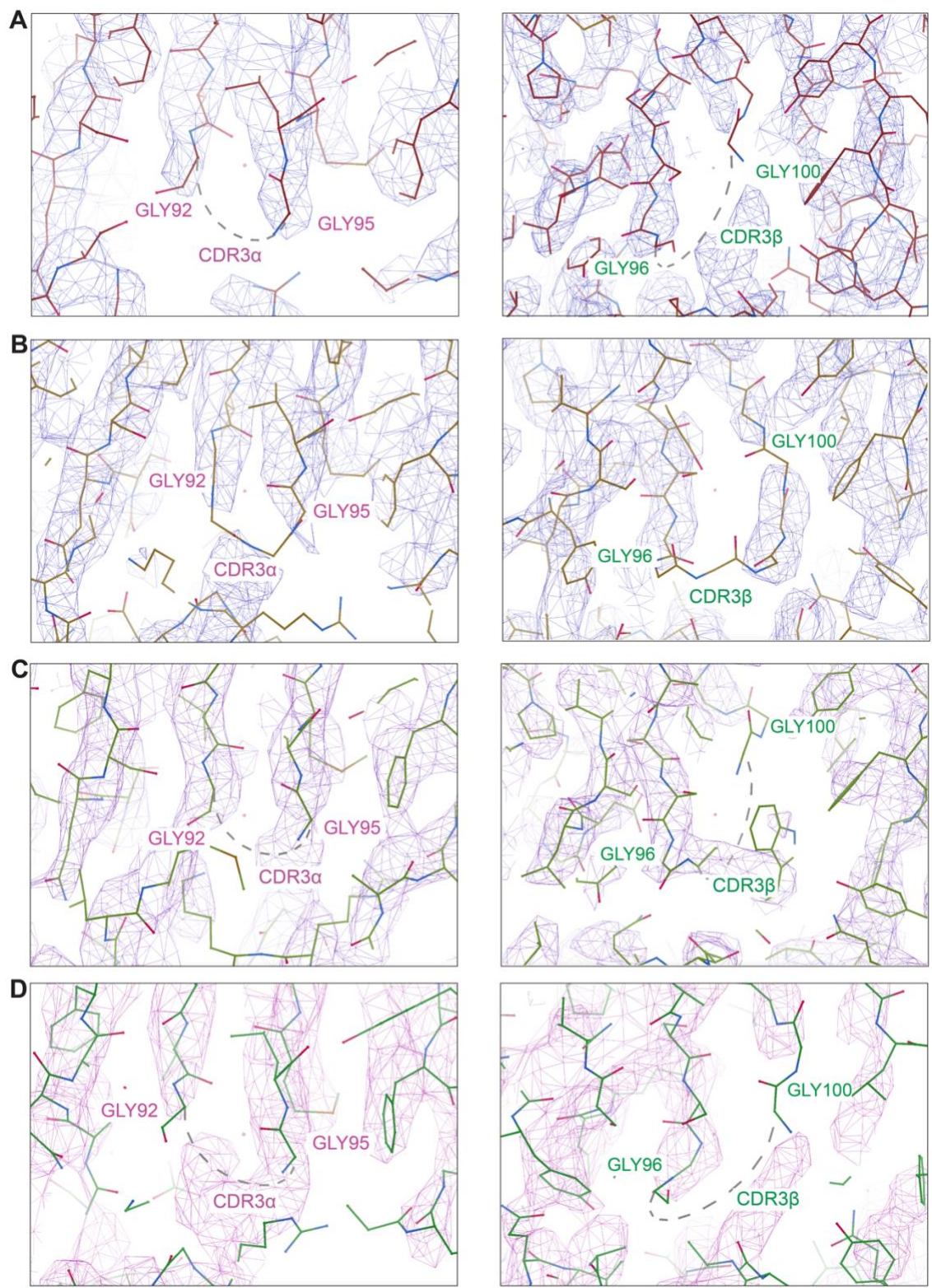

**Figure S6. Cryo-EM density maps showing the absence of CDR3 loop density in germline TCR156-HLA-A\*02 complexes.**

(A-D), Representative cryo-EM density maps (blue mesh) of the CDR3α (left) and CDR3β (right) regions from germline GL156-PAP-A2 (A), GL156-MART1-A2 (B), GL156-NYESO-A2 (C), and GL156-polyA-A2 (D) complexes. In all structures, the engineered poly-glycine CDR3$\alpha/\beta$ loops lack interpretable density (dashed outlines), consistent with high conformational flexibility and minimal engagement with the pMHC surface. Only in the MART1 complex, complete CDR3 loops were sufficiently resolved to permit model building; in the remaining complexes, residues CDR3α Gly93-Gly94 and CDR3β Gly97-Gly99 could not be modeled due to missing density.

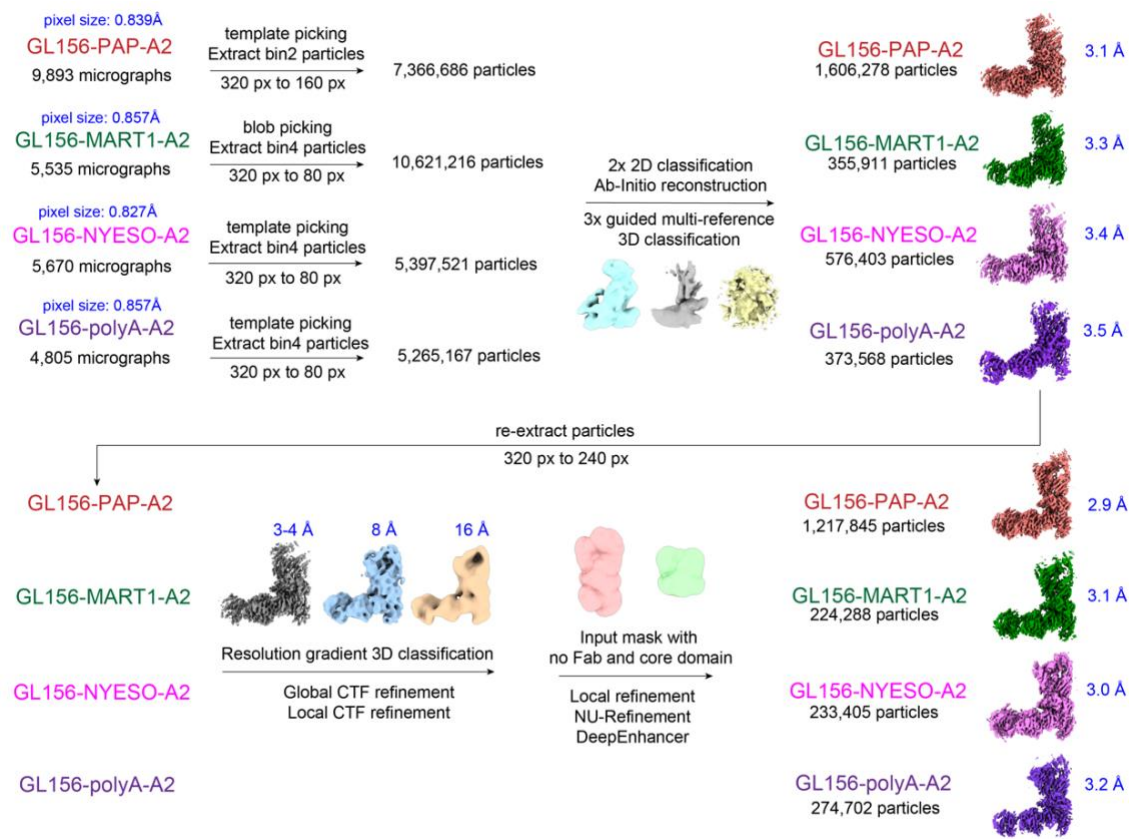

**Figure S7. Cryo-EM data processing workflow for germline TCR156-HLA-A\*02 complexes.**

Overview of particle picking, 2D/3D classification, and refinement procedures for the four germline complexes (GL156-PAP-A2, GL156-MART1-A2, GL156-NYESO-A2, GL156-polyA-A2). Micrograph numbers, particle counts, and intermediate processing steps are indicated for each dataset. Final cryo-EM reconstructions and their corresponding resolutions (Å) are shown on the right.

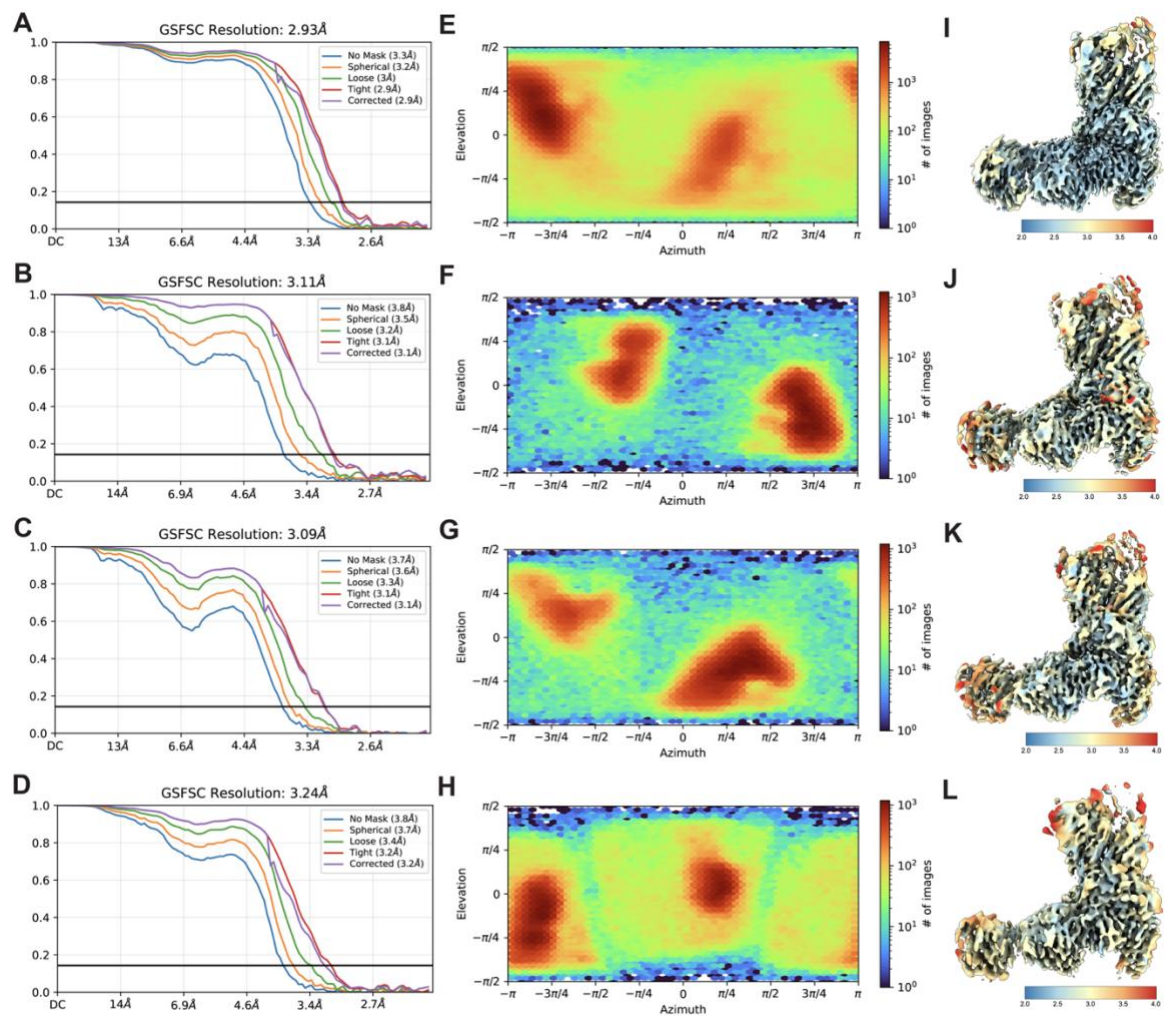

**Figure S8. Global and local resolution assessments of germline TCR156-HLA-A\*02 cryo-EM reconstructions.**

(A-D) Global resolution for (A) GL156-PAP-A2, (B) GL156-MART1-A2, (C) GL156-NYESO-A2, and (D) GL156-polyA-A2. Each panel includes the gold-standard Fourier shell correlation (GS-FSC) curve with the global resolution reported at the 0.143 threshold.

(E-H) Particle orientation analysis for the corresponding complexes in (A-D). Each panel includes an angular distribution heat map illustrating the sampling of particle orientations during 3D refinement.

(I-L) Local resolution maps for the corresponding complexes in (A-D). Density maps are colored according to local resolution estimates calculated in cryoSPARC using an FSC threshold of 0.143. A color bar indicates the resolution gradient in Ångströms (Å), where deep blue represents the most rigid core and red indicates increased conformational flexibility in peripheral domains.

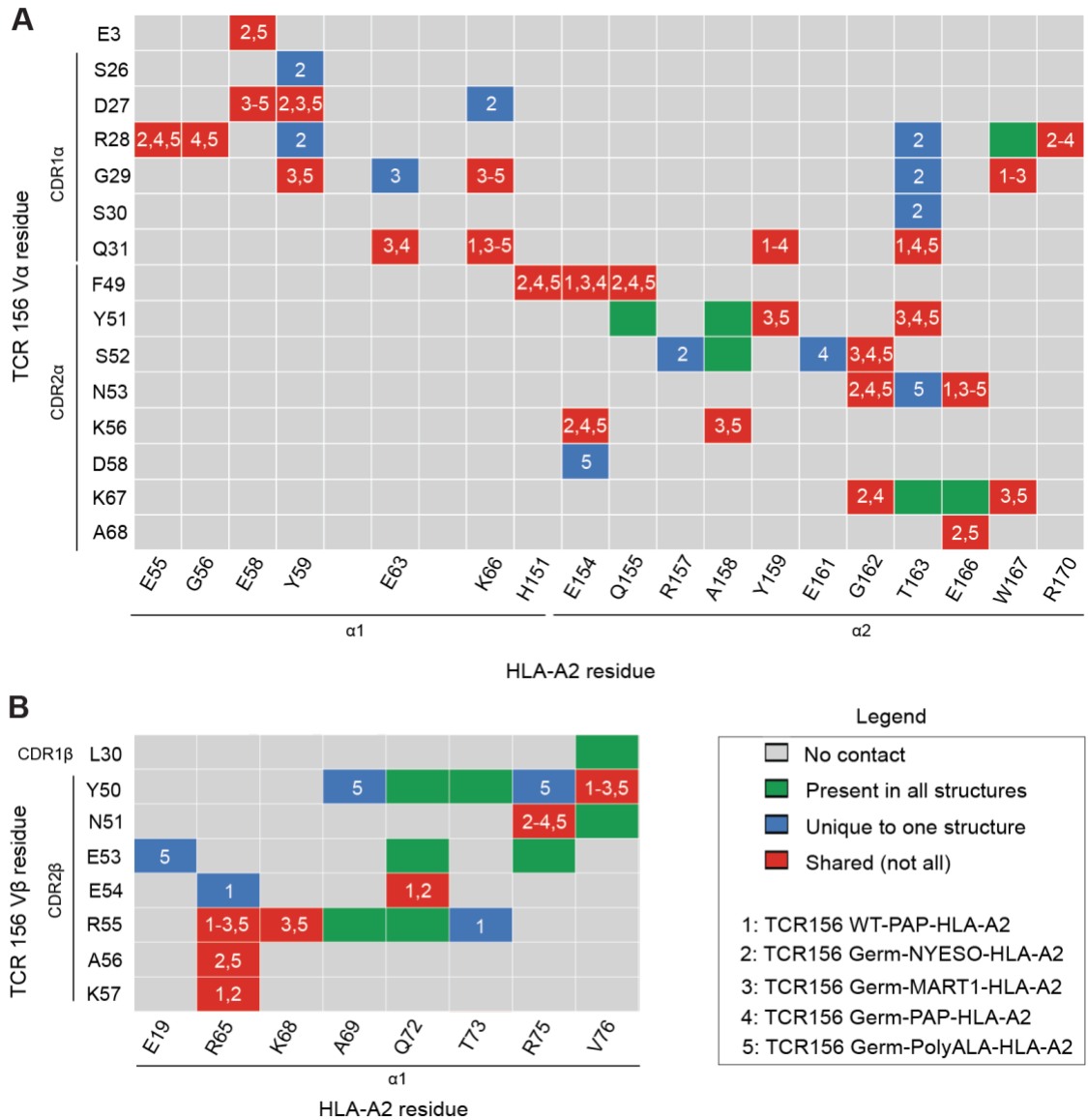

**Figure S9. Germline-like TCR156 CDR1/CDR2 contacts with HLA-A\*02 are conserved across peptide contexts.**

(A-B) Contact maps between (A) TCR156 Vα (CDR1α and CDR2α) or (B) Vβ (CDR1β and CDR2β) residues (rows) and HLA-A02 residues on the $\alpha 1$ and $\alpha 2$ helices (columns) for five structures: Structure 1: WT TCR156-PAP-HLA-A02; 2: germline TCR156-NYESO-HLA-A02; 3: germline TCR156-MART1-HLA-A02; 4: germline TCR156-PAP-HLA-A02; 5: germline TCR156-poly-Ala-HLA-A02. Each square indicates whether a given residue pair makes no contact (gray), a contact present in all five structures (green), a contact unique to one structure (blue), or a contact shared by some but not all structures (red). Contacts were defined using a combination of PDBePISA, PyMol and direct inspection of structures with distances $\leq 3.6$ Å.

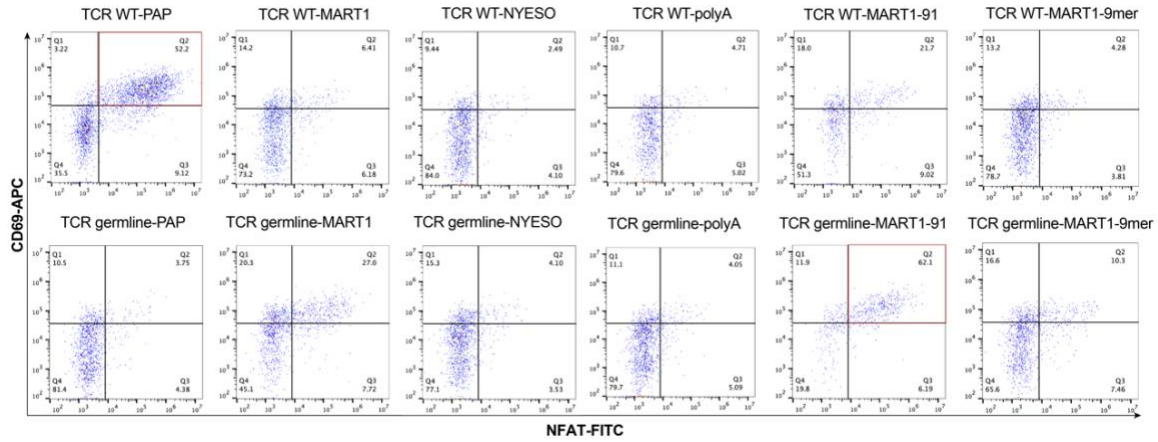

**Figure S10. Functional analysis of WT and germline TCR activation in response to peptide stimulation.**

Representative flow cytometry plots showing CD69-APC versus NFAT-FITC staining for WT and germline TCR156 under each peptide stimulation condition.

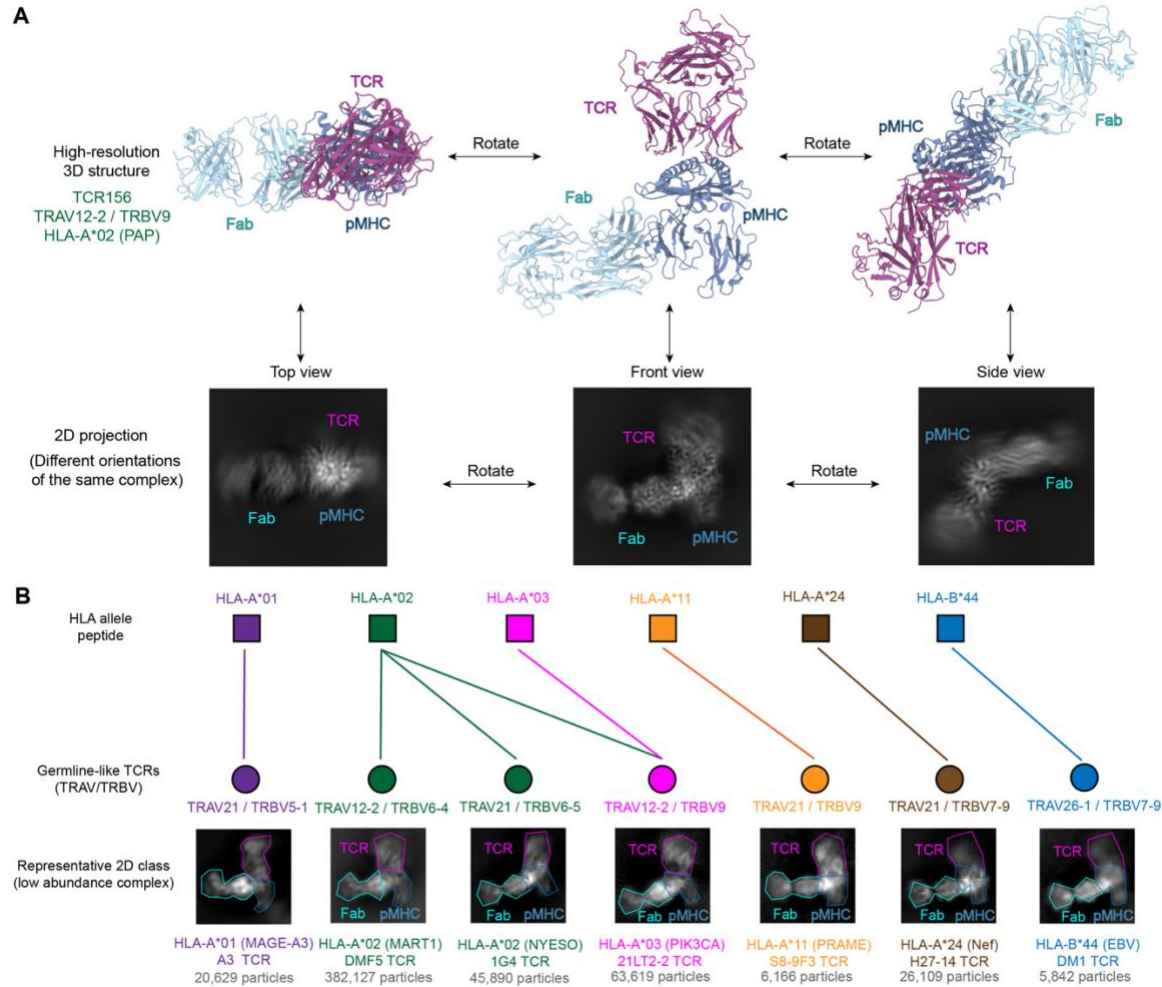

**Figure S11. Germline-mediated docking geometries are conserved across distinct TCR lineages and broadly accessible across HLA alleles.**

(A) Representative 3D cryo-EM structures and their corresponding 2D class averages. Top: Atomic model of the germline-like TCR156–PAP–HLA-A\*02 complex shown in three orthogonal orientations. TCR is shown in magenta, pMHC in blue, and β2M Fab in cyan. Bottom: Representative 2D class averages corresponding to similar viewing orientations of the same complex. The characteristic spatial arrangement of the TCR, pMHC, and Fab domains in the 2D projections enables identification of intact germline-like TCR-pMHC assemblies.

(B) Broad accessibility of germline-like TCR-pMHC complex formation across multiple TCR lineages and HLA alleles. Schematic summary of representative germline-like TCRs generated using distinct TRAV/TRBV combinations and tested against different HLA class I alleles and peptides. Representative 2D class averages are shown below each complex, demonstrating detectable formation of low-abundance germline-like TCR-pMHC particles across multiple HLA backgrounds, including HLA-A\*01, HLA-A\*02, HLA-A\*03, HLA-A\*11, HLA-A\*24, and HLA-B\*44. Particle numbers correspond to selected particles contributing to representative 2D classes.

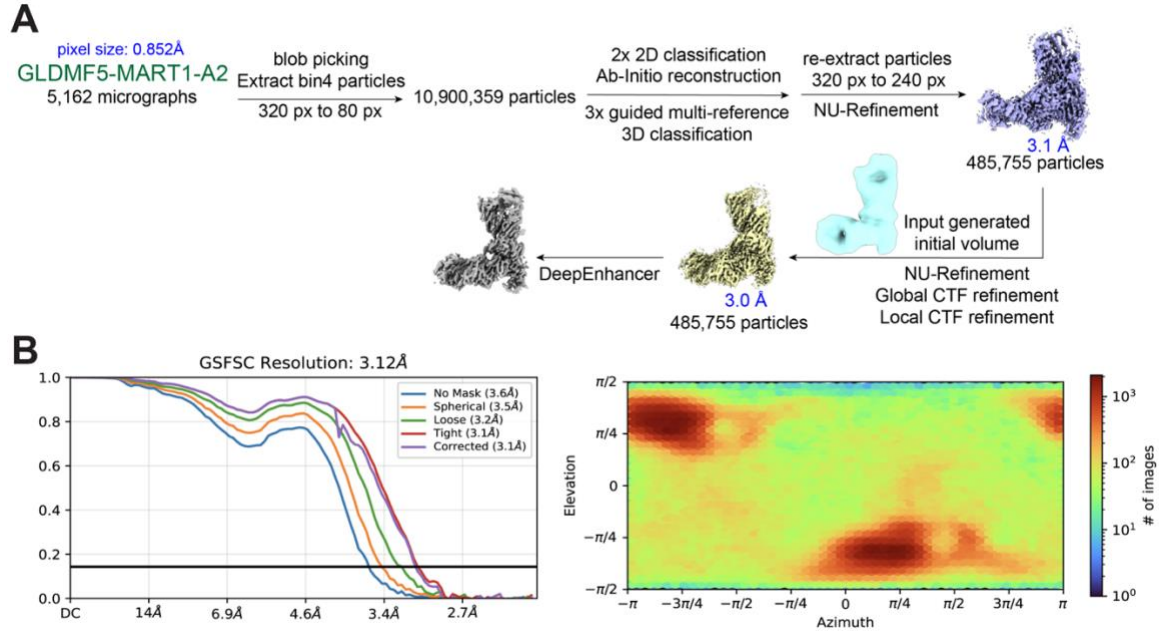

**Figure S12. Cryo-EM data processing workflow for germline DMF5-HLA-A\*02.**

(A) Overview of particle picking, 2D/3D classification, refinement procedures, and particle numbers for GLDMF5-MART1-A2 dataset. Representative intermediate 3D classes, final particle subsets, and the resulting cryo-EM maps with their resolutions (Å) are shown.

(B) Gold-standard Fourier shell correlation (GS-FSC) curves and corresponding angular distribution heat maps for the germline complexes GLDMF5-MART1-A2.

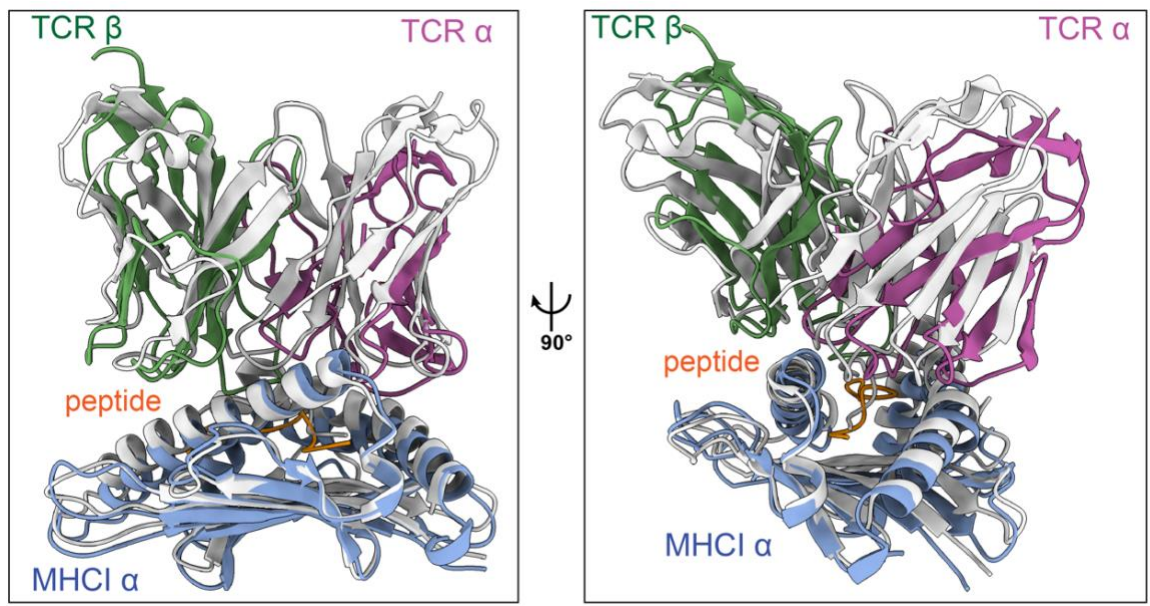

**Figure S13. Distinct docking geometries of TRAV12-2 TCRs on HLA-A\*02.**

Superposition of the WT TCR156-PAP-HLA-A\*02 complex (colored; TCRα in magenta, TCRβ in green, peptide in orange, HLA-A\*02 $\alpha$ chain in blue) with the WT DMF5-MART1-HLA-A\*02 complex (gray), shown in two orthogonal views.

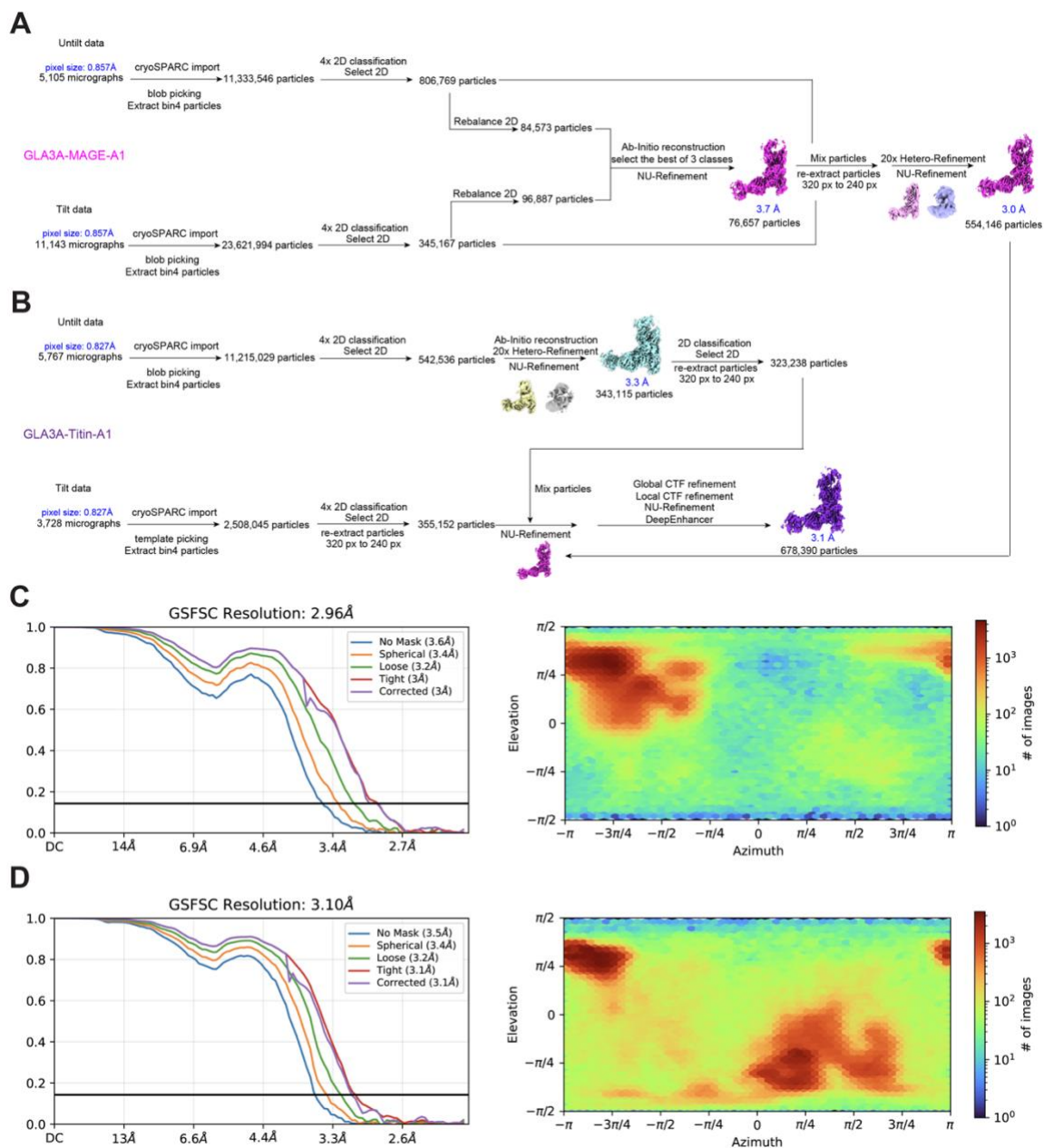

**Figure S14. Cryo-EM data processing workflow for germline A3A-HLA-A\*01 complexes.**

(A-B) Overview of particle picking, 2D/3D classification, refinement procedures, and particle numbers for (A) GLA3A-MAGE-A1, and (B) GLA3A-Titin-A1 datasets. Representative intermediate 3D classes, final particle subsets, and the resulting cryo-EM maps with their resolutions (Å) are shown.

(C-D) Gold-standard Fourier shell correlation (GS-FSC) curves and corresponding angular distribution heat maps for the germline complexes (C) GLA3A-MAGE-A1, and (D) GLA3A-Titin-A1. Reported GS-FSC resolutions are indicated in each panel, and angular heat maps show the distribution of particle orientations contributing to the final refined maps.

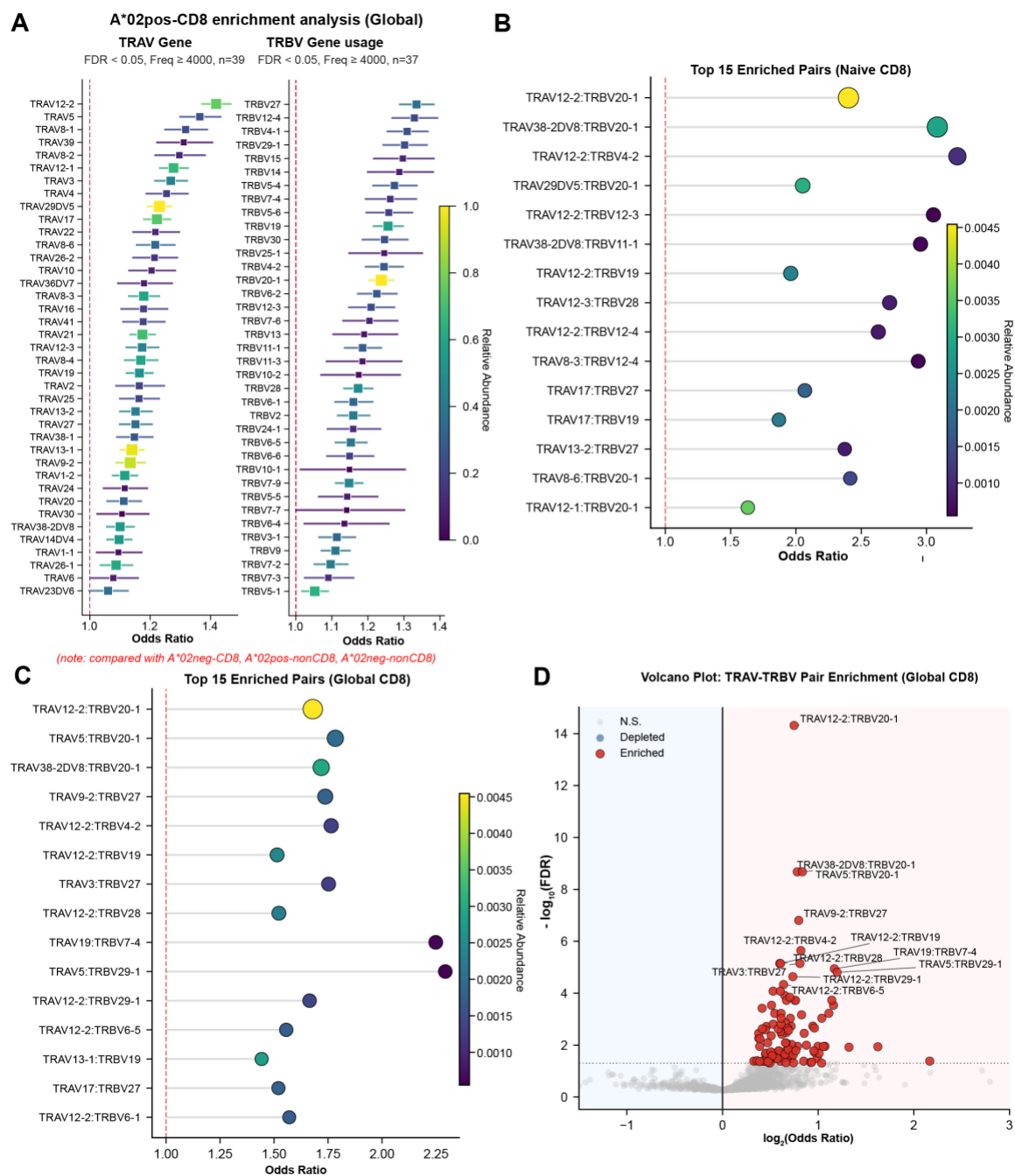

**Figure S15. Expanded analysis of V gene usage and Vα-Vβ pairing enrichment in HLA-A\*02-associated TCR repertoires.**

(A) Enrichment of TRAV and TRBV gene usage in HLA-A02<sup>+</sup> CD8<sup>+</sup> T cell clonotypes. Enrichment was calculated by comparing A02<sup>+</sup> CD8<sup>+</sup> clonotypes against combined control groups (A02<sup>-</sup> CD8<sup>+</sup>, A02<sup>+</sup> non-CD8, and A\*02<sup>-</sup> non-CD8 clonotypes). Points represent odds ratios (OR), with horizontal bars indicating confidence intervals. Only genes meeting significance (FDR < 0.05) and frequency thresholds (≥ 4,000 clonotypes) are shown. The dashed vertical line indicates no enrichment (OR = 1), and color scale reflects proportional abundance within the dataset.

(B-C) Enrichment of TRAV-TRBV gene pairings in HLA-A\*02<sup>+</sup> CD8<sup>+</sup> Tn and global CD8 T cells. Each point represents a Vα-Vβ combination, plotted by odds ratio (x-axis), with color indicating proportional abundance and ordered with FDR values. The dashed vertical line indicates no enrichment (odds ratio = 1).

(D) Volcano plot of TRA-TRBV pairing enrichment in HLA-A\*02<sup>+</sup> global CD8<sup>+</sup> clonotypes. Each point represents a Vα-Vβ combination, plotted by $\log_2(\text{odds ratio})$ and $-\log_{10}(\text{FDR})$. Significantly enriched pairs (FDR < 0.05) are highlighted in red, while non-significant pairs are shown in grey. Selected high-confidence enriched pairs are labeled.

**Table S1. Summary of data collection and model statistics.**

|  | AS4.2-PRPF3-HLA-B*27 | GL156-PAP-A2 | GL156-MART1-A2 | GL156-NYESO-A2 | GL156-polyA-A2 |
| --- | --- | --- | --- | --- | --- |
| **Data collection** |  |  |  |  |  |
| EM equipment | FEI Titan Krios |  |  |  |  |
| Voltage (kV) | 300 | 300 | 300 | 300 | 300 |
| Detector | K3 | K3 | K3 | K3 | K3 |
| Pixel size (Å) | 0.852 | 0.839 | 0.857 | 0.827 | 0.857 |
| Electron dose (e-/Å<sup>2</sup>) | 60 | 60 | 60 | 60 | 60 |
| Defocus range (µm) | 1.0~2.0 | 1.0~2.0 | 1.0~2.0 | 1.0~2.0 | 1.0~2.0 |
| **Reconstruction** |  |  |  |  |  |
| Software | CRYOSPARC |  |  |  |  |
| Number of used Particles | 832,124 | 1,217,845 | 224,288 | 233,405 | 274,702 |
| Map sharpening Method | DeepEMhancer (visualization) |  |  |  |  |
| Final Resolution (Å) | 2.8 | 2.9 | 3.1 | 3.0 | 3.2 |
| **Model building and refinement** |  |  |  |  |  |
| Software | Phenix |  |  |  |  |
| Initial models used (PDB codes) | 7N2N | 9NMU | 9NMU<br>+3QDG | 9NMU<br>+9C3E | 9NMU |
| Model composition |  |  |  |  |  |
| Non-hydrogen atoms | 6681 | 6523 | 6547 | 6533 | 6509 |
| Protein residues | 836 | 815 | 821 | 815 | 815 |
| Ligand | ---- | ---- | ---- | ---- | ---- |
| B factors (Å<sup>2</sup>) |  |  |  |  |  |
| Protein | 66.70 | 81.59 | 89.61 | 80.81 | 99.02 |
| R.m.s deviations |  |  |  |  |  |
| Bonds length (Å) | 0.004 | 0.003 | 0.003 | 0.004 | 0.004 |
| Bonds Angle (°) | 0.917 | 0.506 | 0.511 | 0.562 | 0.569 |
| Ramachandran plot statistics (%) |  |  |  |  |  |
| Preferred | 96.00 | 95.38 | 94.70 | 92.01 | 93.38 |
| Allowed | 4.00 | 4.62 | 5.06 | 7.87 | 6.62 |
| Outlier | 0.00 | 0.00 | 0.25 | 0.12 | 0.00 |
| Validation |  |  |  |  |  |
| Molprobity score | 1.97 | 2.04 | 2.28 | 2.41 | 2.25 |
| Clash score | 15.50 | 16.55 | 24.18 | 22.74 | 21.33 |
| Rotamer outliers (%) | 0.54 | 0.42 | 1.13 | 1.27 | 0.57 |

**Table S2. Summary of data collection and model statistics.**

|  | GLDMF5-MART1-A2 | GLA3A-MAGE-A1 | GLA3A-Titin-A1 |
| --- | --- | --- | --- |
| **Data collection** |  |  |  |
| EM equipment | FEI Titan Krios |  |  |
| Voltage (kV) | 300 | 300 | 300 |
| Detector | K3 | K3 | K3 |
| Pixel size (Å) | 0.852 | 0.857 | 0.827 |
| Electron dose (e-/Å<sup>2</sup>) | 60 | 60 | 60 |
| Defocus range (µm) | 1.0~2.0 | 1.0~2.0 | 1.0~2.0 |
| **Reconstruction** |  |  |  |
| Software | CRYOSPARC |  |  |
| Number of used Particles | 485,755 | 554,146 | 678,390 |
| Map sharpening Method | DeepEMhancer (visualization) |  |  |
| Final Resolution (Å) | 3.0 | 3.0 | 3.1 |
| **Model building and refinement** |  |  |  |
| Software | Phenix |  |  |
| Initial models used (PDB codes) | 3QDG | 5BRZ | 5BS0 |
| Model composition |  |  |  |
| Non-hydrogen atoms | 6396 | 6485 | 6488 |
| Protein residues | 807 | 813 | 813 |
| Ligand | --- | --- | --- |
| B factors (Å<sup>2</sup>) |  |  |  |
| Protein | 85.74 | 83.30 | 89.78 |
| R.m.s deviations |  |  |  |
| Bonds length (Å) | 0.004 | 0.004 | 0.003 |
| Bonds Angle (°) | 0.510 | 0.963 | 0.526 |
| Ramachandran plot statistics (%) |  |  |  |
| Preferred | 95.11 | 93.99 | 94.62 |
| Allowed | 4.89 | 6.01 | 5.26 |
| Outlier | 0.00 | 0.00 | 0.13 |
| Validation |  |  |  |
| Molprobity score | 2.29 | 2.32 | 2.23 |
| Clash score | 18.55 | 20.17 | 14.07 |
| Rotamer outliers (%) | 1.74 | 1.42 | 1.84 |
